# Single-Molecule Nanopore Profiling of p53-TAD’s Conformational Dynamics, Interactions, and Inhibition

**DOI:** 10.64898/2026.08.28.747917

**Authors:** David DeCoeur, Samantha Schultz, Jianhan Chen, Min Chen

## Abstract

Investigating the conformational dynamics of intrinsically disordered proteins (IDPs) is essential to understanding how their structural heterogeneity underlies function and how their dysregulation contributes to diseases. Here, we utilized an MspA nanopore-based approach for studying the conformational dynamics and interactions of IDPs at the single-molecule level. The platform was demonstrated using the intrinsically disordered transactivation domain of tumor suppressor p53 (p53-TAD), one of the important proteins in cancer biology. We showed that MspA can stably capture p53-TAD and resolve up to six distinct current states with frequent interconversions, revealing a rich conformational landscape. The nanopore also detected the effect of a cancer-associated double mutational variant, N29K/N30D. Combining experiments with steered molecular dynamics simulations, we showed that the mutant sampled compact conformational states more frequently than wild type, consistent with previous NMR studies. Importantly, the MspA platform enabled direct monitoring of E3 ligase MDM2 binding to p53-TAD and resolved how this interaction is inhibited by anti-cancer compound epigallocatechin gallate (EGCG). Notably, EGCG stabilizes one of the six states sampled by p53-TAD, providing a mechanistic explanation for its inhibitory effect. Together, these findings demonstrate the promise of the nanopore platform for label-free monitoring of IDP conformational dynamics, modulation, binding and inhibition at single-molecule resolution.

## Introduction

Intrinsically disordered proteins (IDPs) are a class of functional proteins characterized by an inherent lack of stable tertiary structures under physiological conditions^1–3^. Rather than one stable conformation, they sample dynamic, heterogeneous conformational ensembles, which is central for IDPs to fulfill their roles in cellular communication, regulation, transcription, and more^4–8^.Due to the cellular importance of IDPs, dysregulation of IDPs is implicated in many human diseases, such as neurodegenerative disease, diabetes, and cancers^9–11^. One of the prime examples of IDP dysregulation is p53, a transcription factor involved in promoting cell cycle arrest, apoptosis, and DNA damage repair^12–15^. Dysregulation of p53 is exceedingly common in tumorigenesis; TP53 (the gene encoding tumor suppressor protein p53) is mutated in over 50% of all human cancers^16–19^. The full-length human p53 protein (393 residues) is comprised of four main functional regions: an N-terminal transactivation domain (TAD, residues 1–73), a central DNA-binding domain (DBD, residues 102–292), a tetramerization domain (residues 325–356), and a C-terminal regulatory domain (residues 364–393)^20,21^. p53 is heavily regulated via its intrinsically disordered TAD. Under normal cellular conditions, E3 ligase MDM2 binds p53-TAD for ubiquitylation and subsequent degradation via the proteasome. p53-TAD’s residues 18-26 form a helix which binds to the hydrophobic cleft of MDM2’s N-terminal domain^22^. Phosphorylation of p53-TAD in response to stress, such as DNA damage, disrupts the binding of p53-TAD to MDM2, increasing cellular levels of p53-TAD and allowing cell cycle arrest or apoptosis^23–25^. Inhibition of this interaction between p53 and MDM2 represents a promising strategy for anti-tumor therapies^26–29^.

IDPs represent a promising but challenging class of proteins for drug discovery^30–33^, requiring accurate characterization of their conformational dynamics and interactions for effective therapeutics. Nuclear magnetic resonance (NMR) is commonly used to uncover IDPs’ secondary structure, dynamics, and interactions^34,35^. Small angle X-ray scattering (SAXS) is another popular technique utilized to determine overall shape and interactions of IDPs^36,37^. However, current biophysical techniques used for investigating protein conformation are ineffective at probing IDPs’ structural heterogeneity due to their ensemble averaged measurements. There is a crucial need for techniques capable of probing the heterogeneity of IDP conformation, interactions, and inhibition. Single-molecule techniques, such as single-molecule Förster resonance energy transfer (smFRET)^38,39^, high-speed atomic force microscopy (hsAFM)^40^, and most recently field effect transistors (FETs)^41,42^ have emerged as powerful alternatives to solution-based biophysical investigation for IDPs^43^.

Single-molecule nanopore analysis has recently emerged as a powerful label free method for monitoring the conformational dynamics and inhibition of globular proteins^44–50^. Trapping a single protein analyte within the lumen of a nanopore perturbs the ionic current traversing the pore. Monitoring the current modulation over time can reveal the conformation and dynamics of the confined analyte over extended periods with high temporal and structural resolution. For globular proteins, the resulting current fluctuations can often be associated with specific conformational transitions, with these interpretations being supported by structural information and biochemical tools. Applying nanopore analysis to IDPs offers a powerful means of probing their conformational flexibility, although the resulting signals are more challenging to interpret. Nanopore investigations of IDPs have relied largely on short peptide fragments and/or translocation of the chain through the pore to probe interactions and small-molecule effects^51–56^. These approaches restrict what can be resolved, as short peptide fragments cannot report on the long-range contacts and ensemble behavior that define the full-length protein, while translocation reduces the ability to observe natural transitions between conformational states as they occur in real time. Resolving these dynamics instead requires trapping the full-length IDP within the pore’s lumen, as has been successfully applied to globular proteins^44–49^.

Here, we present an MspA nanopore-based platform to investigate the conformational dynamics of full-length p53-TAD (residues 1-73) and show that it can resolve the conformational landscape and effects of a cancer-associated mutant. We further demonstrate that the platform enables direct monitoring of p53-TAD’s interactions with MDM2 and the effects of a small molecule anti-cancer inhibitor. Our results do not only establish protein nanopores as powerful tools for label-free analysis of IDP structure and interaction, but also reveal the conformational landscape of p53-TAD under nanopore confinement and how this landscape is modulated by mutation, MDM2 binding, and small-molecule inhibition.

## Results

### Trapping wild type p53-TAD to resolve conformational substates

Due to the net negative charge of p53-TAD (~ −15.75e at pH 7.4, Fig. S1), we drove capture of the IDP into MspA via electrophoresis from cis to trans (Fig. 1a) at positive applied potentials, causing in a reduction in the open pore current (*I*_*o*_) and characterized by the resulting current signal (*I*) upon capture of the IDP into MspA’s lumen. The average dwell times of the p53-TAD events increased with increasing potentials from +100 mV to +200 mV (Fig. S2), which can be explained by the increased electrophoretic force pulling p53-TAD into MspA and holding it within the lumen, resulting in longer duration capture. In addition, the Stokes radius of p53-TAD is ~2.38 nm^57^ while the constriction site of MspA is ~1.2 nm across^58^. Thus, we concluded that p53-TAD unlikely was translocating through MspA at the voltage range investigated. At +200 mV, p53-TAD induced current blockade events with diverse residual current distributions (Fig. 1b and Fig. S3). These events could sample multiple current states, defined by their normalized residual current level 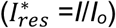 (Fig. 1c-d), ranging from near complete blockage 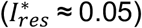 to only slight current blockage 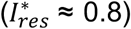. All-points histograms of capture events showed six distinct duration-weighted 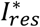 peaks centered at 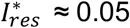, 0.15, 0.28, 0.45, 0.66, and 0.8, denoted as S1, S2, S3, S4, S5, and S6 states, respectively (Fig. S4).

**Figure 1:**
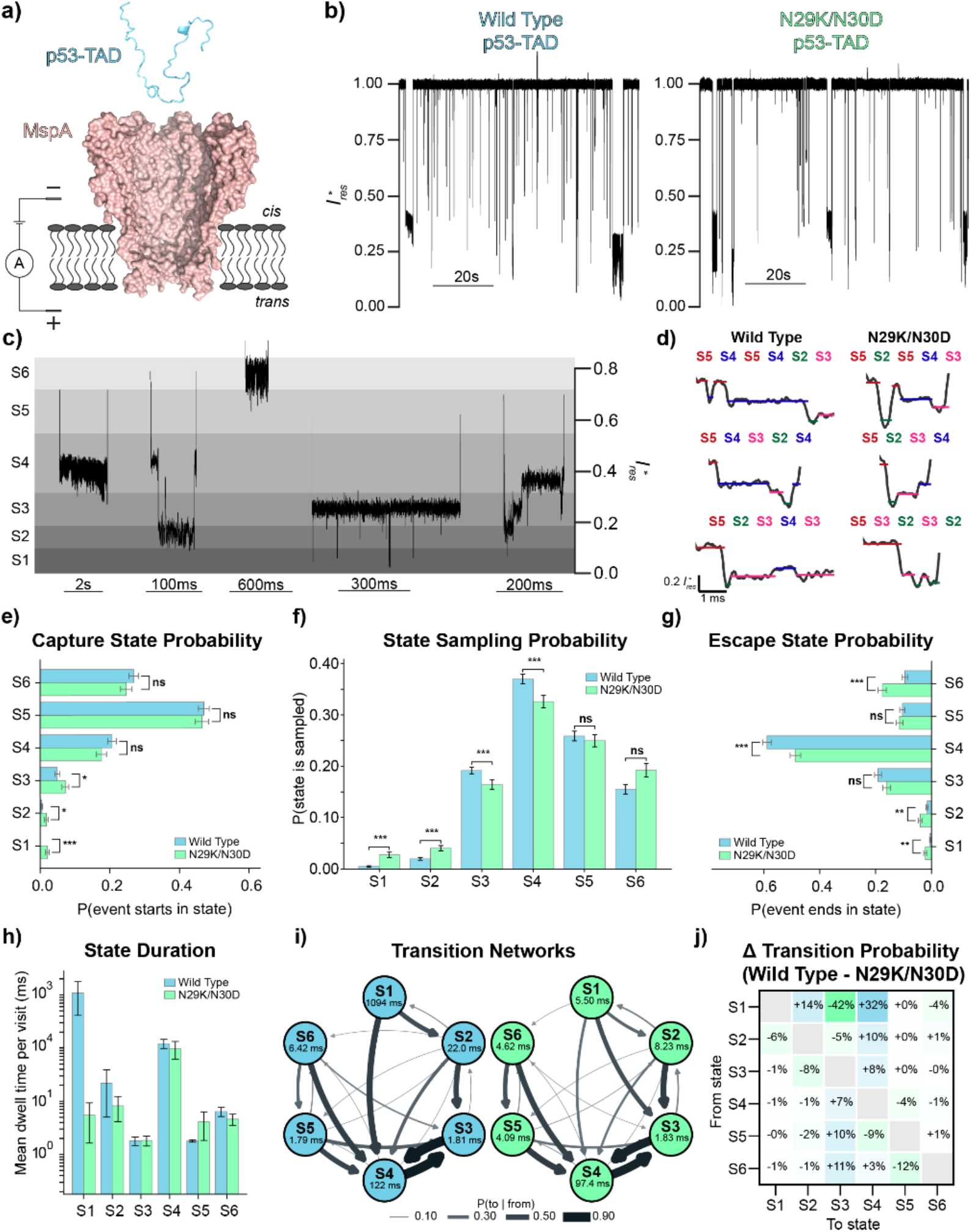
Resolving conformational ensemble shift of cancer-associated mutant p53-TAD. **a)** Cartoon schematic of MspA nanopore (PDB:1UUN^*58*^) analysis with p53-TAD in cis chamber. **b)** Representative current recording traces of wild type and N29K/N30D p53-TAD. Current recordings were acquired in 150mM NaCl, 20mM HEPES, pH 7.4 at +200mV. **c)** Zoomed-in view of individual events of wild type and N29K/N30D. The gradient shading backdrop represents the residual current windows occupied by the observed 6 characteristic states (S1-S6) sampled by p53-TAD variants. **d)** Representative state assignment for wild type and N29K/N30D events shown as raw current drawn in black with the associated state assigned to the time series overlaid in color. State names above events show the sequence of sampled states for each capture event. **e)** Probability of p53-TAD variants being captured in each state. Error bars represent the standard error of mean (ns denotes p-value > 0.05, **\*** denotes p ≤ 0.5, **\*\*** denotes p ≤ 0.01, **\*\*\*** denotes p ≤ 0.001). **f)** State sampling probability of individual wild type and N29K/N30D p53-TAD events. **g)** Probability of p53-TAD exiting in each state. **h)** Average dwell time for all 6 states with error bars represent the standard error of mean. **i)** Transition networks of wild type and N29K/N30D. The probability of exiting one state for another in the transition networks is indicated with arrow thickness (self-transition information encoded in the mean dwell time labeled under each state name). **j)** The difference in transition probability between wild type and N29K/N30D events (wild type – N29K/N30D) with saturation of shading indicating higher likelihood of transitions for one variant (blue = wild type more likely, green = N29K/N30D more likely).

Motivated by the heterogeneity of event state levels observed, we utilized a Hidden Markov model (HMM) to determine the sampling probability of each current state and state-dependent transition preferences. The HMM analysis revealed that p53-TAD was captured almost exclusively through high residual current states, with 47.0% of events initiating in S5 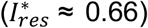 and 26.9% in S6 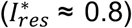 (Fig. 1e). This preference in capture state indicates an initial engagement of p53-TAD with the upper vestibule. In addition, p53-TAD samples medium 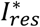 states S3, S4, and S5 most often (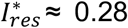, 0.45, 0.66 respectively), with low residual current states S1 and S2 (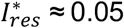 and 0.15 respectively) rarely sampled (Fig. 1f). We interpreted the low 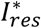 states as compacted conformations of p53-TAD capable of fitting very close to the constriction site within MspA, therefore blocking the majority of the ionic current. Medium current states were interpreted as representing more expanded conformational states that require more space to sample, leading to occupancy of p53-TAD higher in the pore’s vestibule. The high 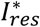 state S6 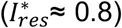 could represent multiple phenomena, such as incomplete capture of the IDP within MspA or potentially transient interactions with the pore that leads to partial blockage of ionic current. p53-TAD preferentially escaped from MspA from S4 (58.9%) with S3 as a secondary exit state (19.1%) (Fig. 1g). This result indicates that p53-TAD preferentially escapes from the nanopore with a different configuration than the configurations it was sampling when captured. We found each p53-TAD triggered event had an average of 1.9 current state transitions to other states (Fig.S5) and the per-state dwell times varied drastically across the six HMM-assigned current states (Fig. 1h). Medium 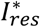 states S3 and S5 were the most transient, with mean dwell times of 1.81 ms and 1.79 ms per visit respectively, while S6 (6.42 ms) and S2 (22.0 ms) occupied intermediate timescales. S4 was the highest sampled state with a relatively long average duration of 121.95 ms per visit. S1 was the longest duration state, with a mean dwell time of 1093.87 ms per visit; however, it was rarely sampled (19 total visits across 1040 events). Analysis of the off-diagonal transition matrix revealed that the conformational dynamics of wild-type p53-TAD within MspA was dominated by a tight S3↔S4 oscillation, with 90% of S3 exits transitioning to S4 and 89% of S4 exits returning to S3, effectively constituting a two-state kinetic trap within the larger six-state system (Fig. 1i and Fig. S5). Transitions from higher 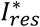 states S5 and S6 were biased toward the S3/S4 basin, with S5 exiting to S4 (48%) and S3 (34%), and S6 exiting to S4 (55%) and S5 (29%). These transitions indicate a general funnel toward mid-pore occupancy.

It is unlikely that each sampled current state represents individual conformations of p53-TAD due to the persistence of each current state lasting much longer than the reconfiguration time of IDPs as well as the degeneracy of possible p53-TAD configurations capable of producing each observed current state. We instead considered that each discrete current state could represent multiple configurations of p53-TAD that have similar properties (such as radius of gyration (*R*_*g*_)) that are being sampled within MspA’s lumen. The transition observed between these states would then represent a transition to an alternative conformational ensemble as confined by the local pore geometry. This interpretation of p53-TAD’s signal in which conformational state determines spatial position within the nanopore would allow us to probe how the conformational ensemble sampled by p53-TAD within the pore may shift upon perturbations such as sequence mutation and cofactor binding.

### Resolving mutational effects on p53-TAD’s conformational dynamics

N29K/N30D p53-TAD is a cancer-associated variant of p53, with previous studies suggesting an overall compaction of the conformational ensemble sampled relative to wild type^59,60^. To test if MspA nanopore could resolve these differences, we analyzed the full length N29K/N30D p53-TAD under the same conditions as wild type. Capture of N29K/N30D p53-TAD within MspA led to events with longer duration than wild type’s with an average of 1.1s ± 5.5 s (compared to 0.46s ± 2.5 s) (Fig. S6). Interestingly, N29K/N30D p53-TAD sampled similar residual current states as wild type (Fig. 1c-d and Fig. S4-8), strengthening our hypothesis that these current states report on the conformational properties of p53-TAD while trapped in MspA. The N29K/N30D mutant exhibited a redistribution of conformational sampling toward deeper pore positions relative to wild type with a higher sampling probability of low 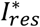 states S1/S2 and reduced sampling of the medium 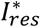 states S3/S4 (Fig. 1f), consistent with the previously reported compaction of the p53-TAD conformational ensemble. Interestingly, despite this increased frequency, the per-visit dwell time of S1 was reduced ~200-fold relative to wild type (5.50 ms vs. 1093.87 ms) (Fig. 1h). The dominant S3↔S4 conformational transition was also observed in N29K/N30D events (82%/82% vs. 90%/89% in wild type), with the transitions redistributed toward lower 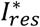 state S2, consistent with expectations (Fig. 1i and Fig. S5). Transitions from S1 were also altered; rather than the jumps to S2 or S4 observed in wild type, the mutant exited S1 preferentially through S3 (42%). Taken together, the enrichment of low 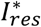 states and reduction of medium 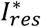 states in N29K/N30D events coupled with previously reported increases in conformational compaction of this mutant relative to wild-type p53-TAD supports that the current states triggered by p53-TAD within MspA can report on the conformational dynamics of the captured IDP.

### Molecular dynamics simulations of p53-TAD in MspA

To further understand the molecular basis of the observed differences between wild-type and N29K/N30D p53-TAD, we employed steered molecular dynamics (SMD) simulations using the hybrid resolution (HyRes) protein force field^61–63^. HyRes represents the protein backbone atomistically while modeling each side chain at intermediate resolution with up to 5 coarse-grained beads, allowing accurate capturing of global and local structural properties of IDPs while substantially reducing the computational cost. Critically, large-scale benchmarking against SAXS, NMR, and smFRET data for over 100 IDPs demonstrates that the latest HyRes model generates dynamic ensembles comparable to, and often more accurate than, those from all-atom simulations in reproducing chain dimensions, transient tertiary contacts, and local secondary structures^63^. We first perform high temperature simulations of wild type and N29K/N30D p53-TAD at 400K for 50 ns to generate ensembles of 300 diverse conformations, which were extracted at evenly spaced intervals from the final 40 ns to avoid bias toward the starting structure (Fig. S9-10). The resulting conformational ensembles of wild type and N29K/N30D p53-TAD had similar compaction to one another (Fig. S11), ensuring the starting conformations would not bias our comparisons between conditions. Each of these individual conformations was then placed 5 nm above MspA for subsequent simulations. To approximate the electrophoretic force driving capture, we screened applied forces of 10, 20, 25, 30, and 45 pN distributed evenly across the negatively charged residues of p53-TAD in the −Z direction toward MspA. The negative residues of p53-TAD are well dispersed throughout the sequence (Fig. S1), allowing relatively even pulling force across the length of the protein. While this method of force application is not a physical representation of the forces involved in protein capture into nanopores (such as the electrophoretic force and its spatial dependence), we intended only to explore how the IDP might capture and sample within MspA differently given the introduced double mutation. Pilot simulations suggested that 10 pN of force across p53-TAD’s negative residues allowed for efficient capture of the protein into MspA while enabling the protein to explore conformational shifts and dynamic interactions with MspA.

We ran replicate SMD simulations for both wild type and N29K/N30D p53-TAD initiated from each of the 300 conformations pre-generated (Fig. S12). By analyzing the displacement of p53-TAD’s center of mass (COM) from the constriction site of MspA across all trajectories, we identified three major positional states shared between the two constructs (Fig. 2a and Fig. S13): a COM position ~50 Å from the constriction, reflecting sampling within the upper vestibule, a preferred position at ~30 Å, representing the global free energy minimum across both conditions, and an additional population ~10 Å from the constriction, directly situated next to the pore’s narrowest point. Comparing the two constructs revealed enrichment of the wild type p53-TAD’s occupancy in the upper area of MspA’s vestibule with the mutant conversely occupying a deeper, more constrained positioning near the constriction site relative to wild type (Fig. 2b), in agreement with the experimentally observed low 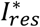 state sampling probability differences. This difference was most strikingly captured in the pseudo potential of mean force (PMF) profiles (Fig. 2c), which displayed a free energy well at ~5 Å for N29K/N30D completely absent for wild type. Averaged across broad positional bins, wild-type p53-TAD experienced a net free energy penalty of ~0.52 *k*_*B*_*T* relative to N29K/N30D within the lower pore region (Z = 0 – 25 Å), while occupying the higher vestibule (Z = 25-75 Å) with an average free energy advantage of ~0.08 *k*_*B*_*T* (Fig. 4d), consistent with the preferential wild type occupancy of the upper pore inferred from nanopore recordings.

**Figure 2:**
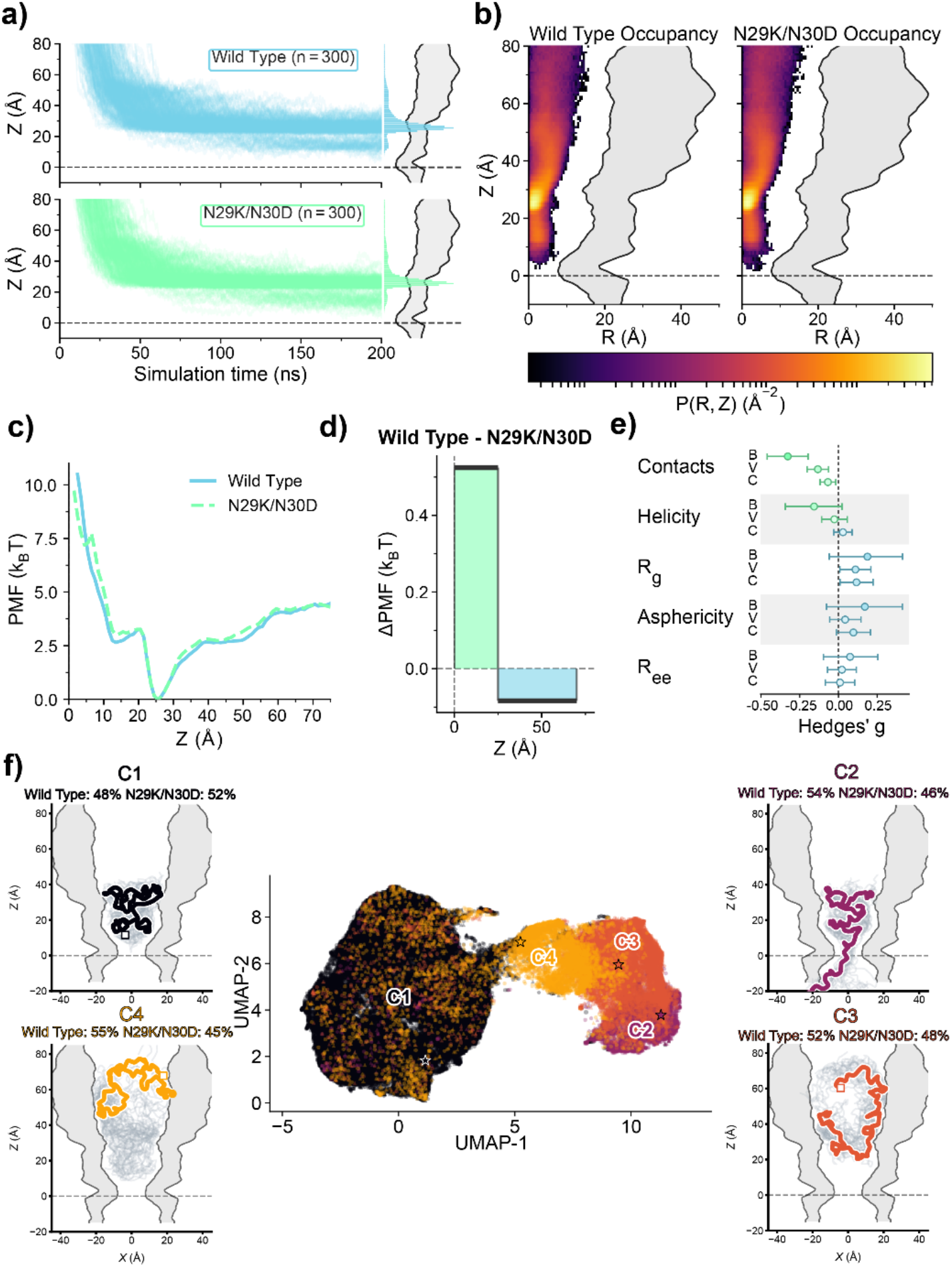
Differences between wild type and N29K/N30D p53-TAD in SMD simulations. **a)** Trajectories of wild type (top, blue) and N29K/N30D (bottom, green) p53-TAD COM distance from MspA’s constriction site (dashed lines at Z = 0 Å) across all 300 simulations, with the distributions shown to the right of each trajectory plot with a longitudinal cross-section of MspA’s inner/outer surface overlaid in grey. **b)** Spatial occupancy heatmaps of wild type (left) and N29K/N30D (right) p53-TAD’s COM within MspA across simulations with MspA’s dimensions overlaid to scale. **c)** Pseudo free energy profiles of both p53-TAD variants as a function of Z axis position within MspA derived from SMD trajectories. **d)** Average free energy difference between wild type and N29K/N30D p53-TAD’s occupancy within the upper (Z > 25 Å) and lower (Z < 25 Å) portions of MspA. **e)** Difference in conformational properties (Contacts: intramolecular contacts) of p53-TAD variants expressed as Hedges’ g (wild type − N29K/N30D; positive values indicate higher in wild type). Bands representations are B: bulk solution ensembles, V: vestibule frames (35-70 Å), C: Near constriction frames (0-30 Å). Error bars represent 95% confidence interval. **f)** Conformational dimensionality reduction and clustering for both variants of p53-TAD by location within MspA. Individual conformations shown as dots and colored by cluster (C1 and C2: Near constriction, C3 and C4: Vestibule). Medoid configurations are indicated with stars and illustrated within MspA as a chain in the cluster color with 25 neighboring conformations from the same cluster overlaid in grey. Variant composition of each cluster is annotated above each representative cluster conformation.

Having established the p53-TAD variants’ preferential occupancy within MspA, we asked whether the variants also differed in conformational properties. We binned configurations of p53-TAD by the distance of the p53-TAD center of mass from the constriction site, 0–30 Å (Near Constriction) and 35–70 Å (Vestibule), while leaving the intervening 30–35 Å unassigned as a buffer against boundary mixing. The clearest difference between variants was in intramolecular contacts as N29K/N30D p53-TAD was enriched relative to wild type in both pore regions, by roughly half a contact in the vestibule and a third of a contact near the constriction (Fig. 2e). The shape descriptors follow the same trend with wild type being slightly more expanded in both regions (Δ*R*g ≈ +0.2–0.3 Å) and marginally more aspherical with a higher end-to-end distance. To verify whether conformational preferences in dilute solution persist within MspA’s lumen, we ran four independent 400 ns simulations of each variant in isolation. We found the conformational differences between variants within MspA mirror the behaviors of the variants’ free ensembles (Fig. S14). In bulk solution, wild type is likewise the more expanded and more aspherical chain and N29K/N30D the more internally contacted. However, confinement compresses the distinction between the two sequences, with all conformational property difference magnitudes being larger in bulk than when residing in MspA. Together these results support that intrinsic differences in IDP properties produce observable effects in nanopore current recordings, mediated by where the chain resides within the pore’s local geometry.

To characterize the types of configurations adopted in each pore region, we embedded both variants’ configurations from both pore regions into a single shared 2D space and partitioned them by clustering within each region separately (Fig. 2f). By clustering a maximum of 2 classes per positional region, we wanted to effectively see the two conformational poles that are sampled by p53-TAD. In both pore regions the separation was driven primarily through chain expansion. In the vestibule, configurations were characterized via either extended but curved (C-shaped) configurations (C3) or by more self-interacting ones of lower radius of gyration (C4). When near the constriction site, p53-TAD sampled either a majority compact state (C1) or an extended state in which one terminus explored past the constriction and contacted trans-presenting residues of MspA (C2). These regional ensembles may relate to the characteristics of p53-TAD that led to the current blockage states S1–S4 in our experimental HMM, though we emphasize that the number of ensembles recovered here reflects the two-region, two-cluster design. Our SMD protocol drives capture of p53-TAD and employs a coarse-grained force field, so the sampled ensembles are those of a driven process rather than of equilibrium capture. Even so, the agreement between simulated and experimentally measured variant occupancy, together with the conformational differences observed here, supports the notion that nanopore analysis can inform on the conformational dynamics and heterogeneity of IDPs.

### Monitoring the complex formation between p53-TAD and MDM2

Having established the ability of MspA to resolve conformational dynamics of p53-TAD and detect mutation induced ensemble shifts, we next investigated whether nanopore analysis could report on p53-TAD’s interaction with its primary negative regulator MDM2 (Fig. 3a). We first characterized MDM2’s N-terminal domain (residues 17–125), which contains the p53-TAD binding cleft. Under +200mV, MDM2 triggered events ranging from 0.05-0.80 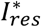 and lasting between 1 – 135 ms long (Fig. 3b and Fig. S16-17). Inclusion of 400 nM p53-TAD in the solution generated new events characterized by near complete blockage of current (~0.05 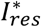, Fig. 3c) with a long duration of over tens of minutes and required manual voltage reversal to eject the trapped proteins from MspA to resume the open pore current. We attributed these new events to the trapping of the p53-TAD/MDM2 complex, as the increased size and charge of the complex relative to either partner alone would be expected to produce lower 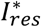 and substantially longer pore residence. Titration of MDM2 to p53-TAD increased the frequency of complex events (calculated via interevent time between events) in a sigmoidal manner (Fig. 3d), yielding an apparent *K*_*d*_ of 308.8 ± 60.4 nM by Hill equation fit. This is consistent with previously reported affinities for the p53-TAD/MDM2 interaction (~240 ± 60 nM ^64,65^. The ability to directly monitor p53-TAD binding via nanopore profiling provides a platform to identify potential small molecule inhibitors of this interaction for therapeutic use.

**Figure 3:**
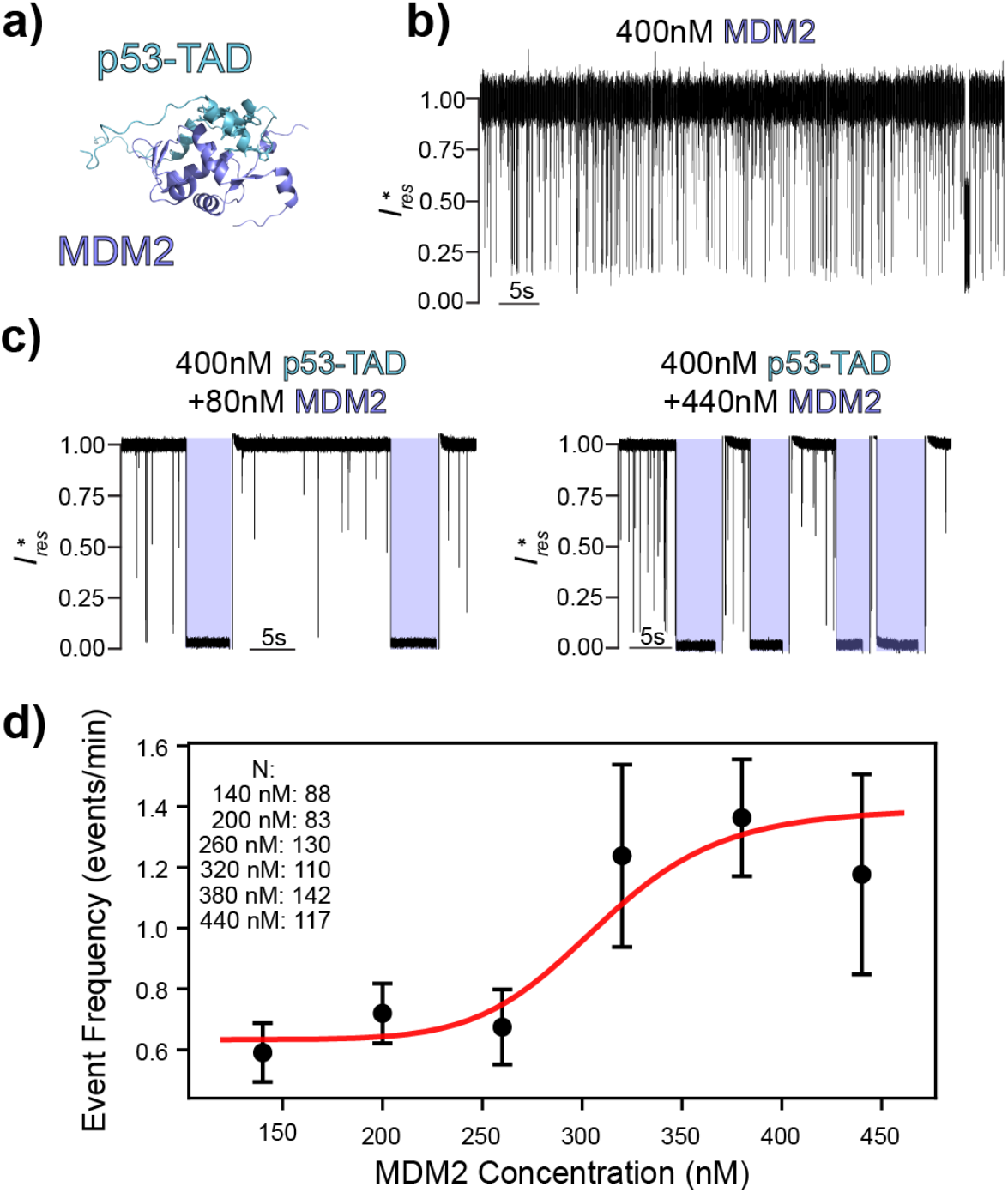
Monitoring complex formation between E3 ligase MDM2 and p53-TAD. **a)** Cartoon depiction of p53-TAD bound to E3 ligase MDM2 (generated via AlphaFold 3^*66*^). p53-TAD residues’ sidechains involved in binding the hydrophobic cleft of MDM2 (residues 18-26) are shown as sticks. **b)** Representative current recording trace of MDM2 alone in 150mM NaCl, 20mM HEPES, 2mM DTT, pH 7.4 solution under +200mV applied potential. **c)** Representative current recording traces of p53-TAD with low and high concentrations of MDM2. The p53-TAD/MDM2 complex capture events are shaded in purple, with each event manually ejected via voltage reversal. **d)** The frequency of p53-TAD/MDM2 complex event plotted as a function of MDM2 concentration with 400 nM p53-TAD. Data points are represented as mean ± SE, with the fitted Hill curve drawn overtop (*K*_d_ = 308.8 nM ± 60.4 nM, R^2^= 0.86).

### Nanopore detection of EGCG binding and inhibition

The ability to directly monitor IDP conformational responses upon small-molecule binding at the single molecule level could present a powerful platform for developing drugs for IDP targets. To validate whether the nanopore system could resolve this conformation modulation, we focused on epigallocatechin gallate (EGCG), the major catechin in green tea that has been to have anti-cancer effects^67,68^. Atomistic simulations together with SAXS previously showed that EGCG induces conformational compaction of p53-TAD to inhibit MDM2 binding^65^ (Fig. 4a). Addition of EGCG alone at +200 mV, did not trigger any current signal change (Fig. 4b). In contrast, when both p53-TAD and EGCG were added, a single long duration event type characterized by ~0.4 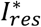 appeared (Fig. 4c). Like the MDM2 bound p53-TAD complex events, EGCG bound p53-TAD rarely left the pore spontaneously, requiring reversing the voltage to eject the drug bound analyte. The residual current level of these events overlapped with the S4 state frequently sampled by apo p53-TAD, though with substantially increased duration. It is likely that EGCG binding stabilizes a subset of p53-TAD conformations corresponding to mid-pore occupancy and restricts the conformational ensemble to states favored at this vestibule depth, reducing the conformational heterogeneity in p53-TAD.

**Figure 4:**
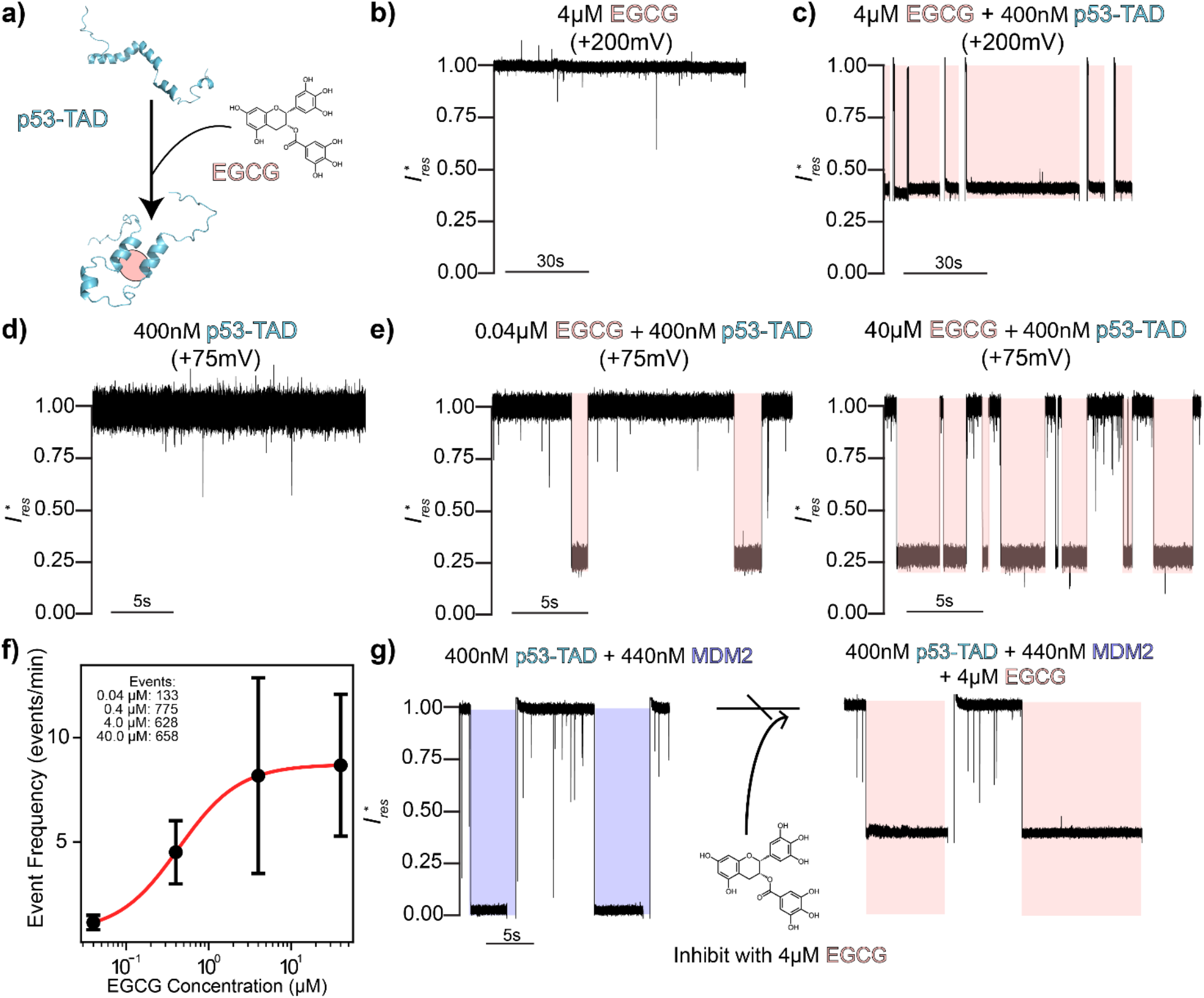
EGCG modulates p53-TAD conformation and inhibits MDM2 binding. **a)** Cartoon illustrating compaction of p53-TAD structure upon binding to EGCG. **b)** Representative current recording trace of EGCG alone in 150mM NaCl, 20mM HEPES, pH 7.4 solution under +200mV applied voltage. **c)** Representative current recording trace of EGCG bound p53-TAD events under +200mV applied voltage (events are ejected via reversal of voltage, followed by return to +200mV). **d)** Representative current recording trace showing a lack of p53-TAD capture under +75mV applied voltage. **e)** Representative current recording traces showing reversible trapping of EGCG bound p53-TAD under low (0.04 µM, left) and high (40 µM, right) EGCG concentrations. **f)** Frequency of EGCG bound p53-TAD events under +75mV applied voltage plotted as a function of EGCG concentration at constant p53-TAD concentration (400nM). Data points represented as mean ± SE, with Hill curve drawn overtop (*K*_d_ = 0.43 µM ± 0.49 µM, *R*^2^= 1.0). **g)** Representative current recording traces of EGCG mediated inhibition of the p53-TAD/MDM2 complex. Characteristic p53-TAD/MDM2 complex capture events (shaded in purple) are ablated upon addition of EGCG to the solution, replaced by EGCG bound p53-TAD events (shaded in pink).

Lowering the applied potential to +75mV reduced the duration of EGCG-bound p53-TAD events. At this voltage, no events were observed with p53-TAD alone (Fig. 4d). However, dosing EGCG to p53-TAD resulted in events with ~0.25 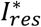 (Fig. 4e). The selective capture of EGCG bound p53-TAD but not apo p53-TAD at +75 mV is interesting and likely multifactorial. First, EGCG binding is expected to reduce the entropic penalty of confining the dynamic IDP within the nanopore, lowering the electrophoretic force required for capture and trapping inside the pore. Additionally, EGCG has a p*K*_*a*_ of ~7.68^69,70^, allowing some population of negatively charged species bound to p53-TAD (pH 7.4), again increasing the electrophoretic force of capture. Interestingly, the residual current levels of EGCG-bound p53-TAD at +75 mV is different from it at +200 mV (~0.25 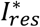 vs ~0.4 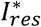 respectively). We hypothesize the lower 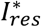 state under +75 mV potentially represents preferential capture of a distinct subset of EGCG-bound p53-TAD than is captured under +200mV, leading to deeper pore occupancy; however underlying mechanism behind this phenomenon is unknown. Titrating EGCG at +75 mV to p53-TAD produced a dose-dependent increase in event frequency (Fig. 4f). Fitting the concentration-dependent frequency to a Hill curve yielded an apparent *K*_*d*_ of 0.43 ± 0.49 µM for the EGCG/p53-TAD interaction. This value is approximately an order magnitude lower than the previously reported bulk *K*_*d*_ of 4 ± 2 μM^65^. This discrepancy likely arises because MspA confinement promotes p53-TAD conformational compaction to prepay the entropic cost of EGCG binding (Fig. 4a), further supporting the promise of conformational modulation in targeting IDPs^30,71,72^.

Strikingly, addition of 4 µM EGCG to the solution completely abolished the p53-TAD/MDM2 complex events, with exclusively EGCG-bound p53-TAD events observed (Fig. 4g). This observation can be explained by EGCG-induced conformational changes in p53-TAD disrupting its interaction with MDM2, in agreement with previously reported inhibition of the p53-TAD/MDM2 interaction by EGCG^65^. We note that the bulk solution contains an equilibrium mixture of p53-TAD bound to both EGCG and MDM2, because of comparable affinities of these two binders. It is apparent that the nanopore assay selectively captures species, further enriching for EGCG-bound p53-TAD over the MDM2-bound complex under these conditions. Nonetheless, the results clearly establish the nanopore as an effective platform to screen for inhibitors of specific IDP binding.

## Discussion

Here we demonstrated the MspA’s capability to capture single p53-TAD proteins and resolve the heterogeneity of its conformational dynamics within a confined space. Characterizing these dynamics with a hidden Markov model, we constructed a conformational landscape from the current blockage levels sampled within MspA and showed how that landscape is reshaped by cancer-associated mutation and by small-molecule binding (Fig. 5a). SMD simulations revealed the differential positional occupancy of wild type and N29K/N30D p53-TAD within MspA as well as the conformational differences while trapped, strengthening our interpretation of the resolved current blockage states. Using the configurations sampled within MspA, the simulations also suggested which conformational properties of p53-TAD could give rise to the experimentally observed blockage levels. MspA further reported the binding of MDM2 to p53-TAD at the level of individual complexes and the disruption of that complex by EGCG (Fig. 5b), potentially providing a platform for interrogating inhibitors of IDP-partner interactions through their modulation of the disordered partner’s conformational ensemble.

**Figure 5:**
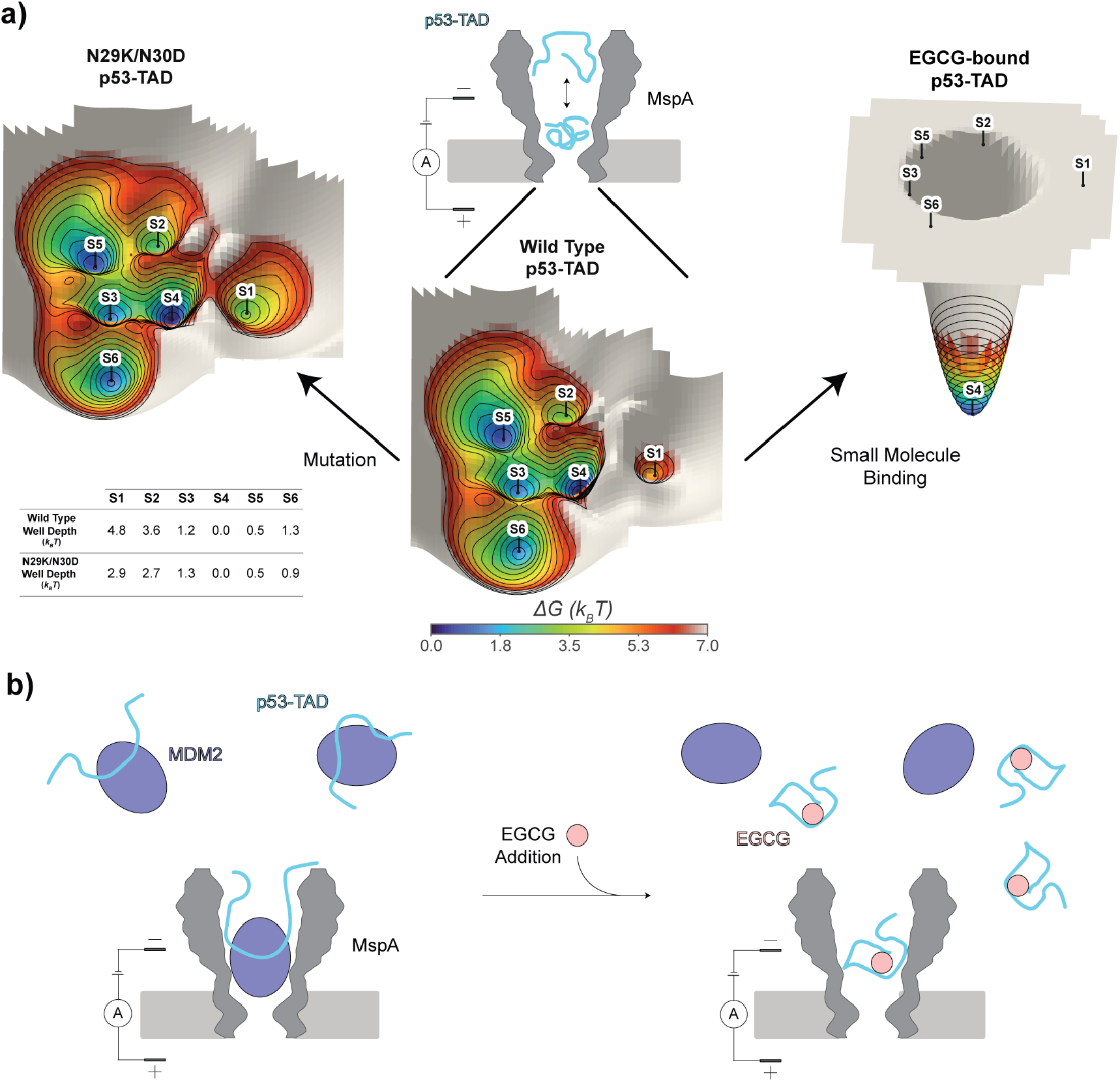
Schematic of MspA’s monitoring of p53-TAD conformational dynamics, modulation, interaction, and inhibition. **a)** Wild type p53-TAD’s conformational landscape within MspA and its reshaping upon either mutational or small molecule perturbations. Both axes are unitless kinetic coordinates from multidimensional scaling of commute times on the decoded transition network with proximity representing kinetic exchange between adjacent states. **b)** Schematic of MspA monitoring the association between MDM2 and p53-TAD. Following drug addition, EGCG modulates p53-TAD’s conformational ensemble, resulting in the inhibition of the p53-TAD/MDM2 complex formation.

Several features of nanopore analysis restrict its interpretation. Firstly, the landscape we report is sampled under confinement within MspA and under the electric field experienced inside the pore, thus the conformational landscape constructed is inherently biased relative to that of the free IDP in solution. Vestibular dimensions represents one facet of this bias, as while MspA’s vestibule comfortably accommodates the reported Stokes radius of p53-TAD (~2.38nm^57^), that property is an ensemble average over a broad distribution of conformations. Extended substates of this distribution are likely excluded sterically, reducing the resolvable conformational space. This limitation could be addressed with the use of a nanopore with a larger internal lumen, such as ClyA^73^, at the cost of smaller fractional blockage (and reduced sensitivity) and with the added requirement that the IDP be conjugated within the lumen to prevent translocation. Nevertheless, the effects of perturbation (both mutational and pharmacological) remain clearly resolvable with the MspA platform, as demonstrated throughout this work. The study of IDPs in their natural cellular environment remains an important long-term goal, as even current industry gold standard in vitro studies of label-free IDPs remain far removed from the crowded, heterogeneous environment in which they naturally function.

In summary, we have demonstrated that single-molecule nanopore analysis can resolve conformational differences between wild-type and disease-associated mutant IDPs, monitor IDP-protein interactions at the single-complex level, detect small-molecule-induced conformational modulation, and report on the inhibition of IDP-partner complexes in real time. These capabilities, combined with the label-free and single-molecule nature of the platform, position nanopore-based approaches as a compelling tool for investigating how conformational heterogeneity mediates IDP function and for identifying therapeutic strategies that target IDPs in the context of human disease.

## Acknowledgments

This project was supported by the US National Institutes of Health grant R01AI156187 (to M.C. and J.C.), R01GM159415 (to M.C.) and R35GM144045 (to J.C.). D.D. was partly supported by the National Institutes of Health T32 GM139789 grant. We thank Dr. Michal Zolkiewski at the Kansas State University for the generous gifts of p53-TAD and MDM2 plasmids. The computational resources for this work were provided by the University of Massachusetts Amherst’s partnership with the Unity Research Computing Platform, a multi-institutional cluster led by the University of Massachusetts and the University of Rhode Island.

## Author Contributions

M.C. and J.C. conceptualized the project. D.D. and S.S. performed the experiments. D.D. performed all simulation and computational analysis. D.D., M.C., and J.C. wrote the manuscript. All authors edited the manuscript. M.C. and J.C. supervised the project.

## Additional information

Extended data is available for this paper at https://doi.org/xxxx

## Supplementary information

The online version contains supplementary material available at https://doi.org/xxxx

