## Supporting information for "Single-Molecule Nanopore Profiling of p53-TAD’s Conformational Dynamics, Interactions, and Inhibition"

**Table of Contents:**

Methods and Materials

Table S1: Simulated conformational properties of p53.

Figure S1: p53-TAD primary sequence annotated.

Figure S2: Duration of wild type p53-TAD voltage study events.

Figure S3: Representative long current trace of wild type p53-TAD capture under +200mV.

Figure S4: All points histograms of wild type and N29K/N30D +200mV events.

Figure S5: State transitions per event and transition matrices.

Figure S6: Duration of wild type and N29K/N30D p53-TAD +200mV events.

Figure S7: Representative long current trace of N29K/N30D p53-TAD capture under +200mV.

Figure S8: Blockage vs duration scatterplots of wild type and N29K/N30D p53-TAD capture events.

Figure S9: Indicative starting conformations of wild type p53-TAD for SMD.

Figure S10: Indicative starting conformations of N29K/N30D p53-TAD

Figure S11: Radius of gyration of starting SMD conformational ensembles.

Figure S12: Simulation system for SMD.

Figure S13: Representative snapshots of p53-TAD at occupied Z positions.

Figure S14: Mutational effect on p53-TAD conformation is preserved under confinement.

Figure S15: Top contact differences between WT and N29K/N30D p53-TAD with MspA.

Figure S16: Representative long current trace of MDM2 Control under +200mV.

Figure S17: $I_{res}^{*}$ vs dwell time for MDM2 control events.

**Materials and Methods**

**Expression and purification of MspA**

M2-MspA-NNN (MspA) protein was expressed and purified as described previously^1^. Briefly, a pT7 plasmid containing MspA with a C-terminal 6x His tag was transformed into BL21(DE3) pLysS competent *E. coli* cells via electroporation and plated on LB agar supplemented with 200 μg/mL ampicillin. A single colony was selected to inoculate 25mL of LB media supplemented with 200 μg/mL ampicillin and incubated overnight at 37°C/300 rpm. After overnight growth, the primary expression culture was diluted to an OD_600_ of 0.075 in LB media supplemented with 200 μg/mL ampicillin to a final volume of 1L and grown at 30°C/300 rpm until it reached an OD_600_ of 0.8. Expression culture was then induced via 0.5 mM IPTG and left to incubate overnight at 16°C/250 rpm. Cells were then pelleted by centrifugation and resuspended in lysis buffer (150 mM NaCl, 50 mM Tris-HCl, pH 8.0, and 0.1 mM PMSF) before lysis (Misonix instrument, 1/2inch probe, 30% amplitude, 2 s pulse, 4 s rest; cycle repeated until OD_600_ of resuspended cells reached ≤ 80% starting value). After lysis, cell membranes were pelleted via centrifugation at 13,000 rpm for 20 min at 4°C. MspA was solubilized by adding 30 mL solubilization buffer (100 mM Na_2_HPO_4_/NaH_2_PO_4_, 0.1 mM EDTA, 150 mM NaCl, 0.5% (v/v) Genapol X–80, pH 6.5) and stirring for 30 min. This sample was then centrifuged again at 13,000 rpm for 30 min at 4°C. Lysate was filtered through 0.22 μm cellulose filter and applied to equilibrated Ni-NTA gravity column (3mL volume, HisPur, ThermoFisher Scientific). Column was washed with 0.5 M NaCl, 20mM HEPES, 0.5% (v/v) Genapol X–80, pH 8.0, followed by 50 mM imidazole, 0.5 M NaCl, 20 mM HEPES, 0.5% (v/v) Genapol X–80, pH 8.0, and eluted with 333 mM imidazole, 0.5 M NaCl, 20 mM HEPES, 0.5% (v/v) Genapol X–80, pH 8.0. MspA containing elution fractions were loaded on a 8% SDS-PAGE gel and ran at 180 V for 60 min. The band corresponding to MspA octamers was then excised from the gel, homogenized with 600 µL of gel extraction buffer (50 mM TrisHCl, 150 mM NaCl, 0.5% Genapol X – 80, pH 7.5) and left on a 60 °C heat block for 30 min. The mixture was then shaken at 50 rpm for 3 hours at 23 °C. MspA oligomer separated via centrifugation at 14000 rpm for 30 minutes at 4°C, aliquoted and stored at -80°C before use.

**Expression and purification of p53-TAD**

Purification of p53-TAD protein variants was performed as described previously^2^. pET28a vectors containing 6xHis-tagged wild type (residues 1-73) or N29K/N30D p53-TAD (gift from Dr. Michal Zolkiewski, Kansas State University) were transformed into BL21(DE3) low background strain cells separately. A single colony of transformed cells was used to inoculate 25mL of LB media supplemented with 50 µg/mL kanamycin and incubated overnight at 37°C/200 rpm. After overnight growth, the primary expression culture was diluted to an OD_600_ of 0.075 in LB media supplemented with 50 µg/mL kanamycin to a final volume of 1L and grown at 30°C/300 rpm until it reached an OD_600_ of 0.8. Expression culture was then induced via 1 mM IPTG and left to incubate overnight at 16°C/200 rpm. Cells were then pelleted by centrifugation and resuspended in lysis buffer (150 mM NaCl, 50 mM Tris HCl pH 8.0, 0.5 mM EDTA, 0.1 mM PMSF) before lysis (Misonix instrument, ½ inch probe, 30% amplitude, 2 s pulse, 4 s rest; cycle repeated until OD_600_ of resuspended cells reached ≤ 80% starting value). Following lysis, supernatant was collected after centrifuging at 13,000 rpm for 30mins at 4°C. Supernatant was collected and applied to equilibrated Ni-NTA gravity column and left to shake on ice for 30min to bind. Ni-NTA column was then washed with 50 mM Tris HCl, 150 mM NaCl, 10 mM Imidazole, pH 8.0, followed by elution with 50 mM Tris HCl, 150 mM NaCl, 250 mM Imidazole, pH 8.0. Eluted fractions were dialyzed overnight in 150 mM NaCl, 50 mM Tris-HCl (pH 8.0), and 5% glycerol with thrombin protease to cleave the 6x His tag. p53-TAD was subsequently purified to remove the thrombin protease via size-exclusion chromatography (Superdex 75 Increase size-exclusion chromatography column, Cytiva Life Sciences) and stored at -80°C.

**Expression and purification of MDM2**

A pGEX-6p-2 vector containing GST tagged MDM2 (residues 17-125) (gift from Dr. Michal Zolkiewski, Kansas State University) was transformed into BL21 (DE3) Rosetta cells. Transformed cells were used to inoculate 25mL of LB media supplemented with 25 µg/mL chloramphenicol and 100 µg/mL of ampicillin and incubated overnight at 37°C/200 rpm. After overnight growth, the primary expression culture was diluted to an OD_600_ of 0.075 in LB media supplemented with 25 µg/mL chloramphenicol and 100 µg/mL of ampicillin to a final volume of 1L and grown at 30°C/300 rpm until it reached an OD_600_ of 0.8. Expression culture was then induced via 1 mM IPTG and left to incubate overnight at 16°C/200 rpm. Cells were then pelleted by centrifugation and resuspended in lysis buffer (150 mM NaCl, 50 mM Tris HCl pH 8.0, 0.5 mM EDTA, 0.1 mM PMSF) before lysis (Misonix instrument, 1/2inch probe, 30% amplitude, 2 s pulse, 4 s rest; cycle repeated until OD_600_ of resuspended cells reached ≤ 80% starting value). Following lysis, supernatant was collected after centrifuging at 13,000 rpm for 30mins at 4°C. Lysate was applied to an equilibrated glutathione Sepharose 4B resin (Cytiva Life Sciences) gravity column and incubated on ice for 60mins. Glutathione column was then washed with 50 mM Tris-HCl, 300 mM NaCl, 2.5 mM EDTA, 2 mM DTT, pH 7.4, then incubated with HRV 3C protease in 50 mM Tris-HCl, pH 7.0, 150 mM NaCl, 1 mM EDTA, 1 mM dithiothreitol overnight at 4°C to cleave MDM2 from column. MDM2 was then collected from column and separated from protease via size-exclusion chromatography (Superdex 75 column, Cytiva Life Sciences) and stored at -80°C.

**Single-channel current recording**

The single-channel recordings were performed in a homemade flow cell containing cis/trans chambers separated by a polytetrafluoroethylene film (thickness of 25 μm, Goodfellow) that has an aperture of ~ 100 μm in diameter at 22 °C. Lipid bilayers are formed by applying 0.5 µL 10% hexadecane (v/v in pentane) to cis/trans sides of the aperture, after ~5min of evaporation followed by addition of 600 µL of chamber buffer (150mM NaCl, 20mM HEPES, pH 7.4), followed by addition of 18 µL of DPhPC in pentane to both chambers and pipetting solution up/down from cis/trans chambers. Ion flow through the nanopore was monitored in voltage-clamp mode with an integrated patch clamp amplifier (Axopatch 200B, Molecular Devices). The current signal was acquired by an analog-to-digital converter (Digidata 1440A, Molecular Devices) at a sampling rate of 50 kHz (for the wild type and N29K/N30D p53-TAD apo characterization experiments) or 10 kHz (for all other experiments) after processing with a four-pole low-pass Bessel filter (2 kHz). After addition of MspA into cis chamber (grounded), it was induced to insert into the lipid bilayer via application of +300 mV potential. All analytes (p53-TAD/MDM2/EGCG) were added to the cis chamber for experiments.

**Nanopore data analysis**

Nanopore recordings were read with pyABF^3^ and analyzed with custom in-house python scripts. Blockade events were extracted from every ABF file using a common threshold-crossing routine. The open-pore current (*I*₀) was determined for each recording, and contiguous samples falling between 0.05 × *I*₀ and 0.80 × *I*₀ were flagged as candidate blockades. Adjacent candidate segments separated by ≤ 1 ms were merged to prevent single blockades from being segmented into multiple events with merged segments shorter than 1 ms were discarded. For each surviving event we recorded start and end times, dwell time, mean current, the root-mean-square deviation of the current about its mean, and the normalized residual current $I_{res}^{*}$ = *I*/*I*₀. All events were manually inspected for verification. Event durations were pooled across all recordings within a condition. Group differences were assessed by two-sided Mann–Whitney *U* tests; for comparisons across more than two groups (e.g. the WT voltage study), an omnibus Kruskal–Wallis test preceded pairwise testing, and *p*-values were adjusted for multiple comparisons by the Bonferroni method (α = 0.05). Event frequency was computed as the inverse of the mean inter-event time between consecutive events (reported in events min⁻¹). For each analyte concentration, frequencies were averaged across replicate recordings and reported as mean ± SEM.

p53-TAD current levels were resolved with a hidden Markov model with Gaussian emissions. Each extracted event trace was low-pass filtered at 10 kHz with a fourth-order Butterworth filter applied in the forward–reverse direction (zero phase distortion), and 0.1 ms was trimmed from each end to remove open-pore transients at the capture and escape edges. Models were parameterized by the number of hidden states, the emission covariance type, and a strong self-transition prior on the diagonal of the transition matrix. Emission means were seeded at approximate $I_{res}^{*}$ levels observed in the all points histograms and held fixed during expectation–maximization, with per-state variances assigned analytically from the Viterbi-decoded sample assignments after fitting and constrained to 10⁻³ ≤ σ² ≤ 0.02 to prevent state collapse or over-broadening. A grid over the number of states and the self-transition prior was evaluated by log-likelihood, AIC, and BIC, with visual inspection of decoded traces used as a sanity check, after which a six-state model was selected. To ensure that state definitions were identical across conditions, a single shared HMM was fit on the pooled wild-type and N29K/N30D data and then used to decode every event by the Viterbi algorithm. Decoded segments shorter than 0.2 ms (10 samples at 50 kHz) were absorbed into whichever neighboring segment had the closest emission mean. Transition matrices are reported in off-diagonal–normalized form, *P*(next = j | current = i, a jump occurs), which isolates branching preference from state stickiness. Occupancy and sampling probability were compared between conditions by two-sided Mann–Whitney *U* tests while categorical capture/escape distributions were compared per-state by Fisher's exact test.

The free energy surfaces were constructed from the Viterbi-decoded state sequences of the shared reference. For each condition we measured how often each state was occupied, averaging across events so that every event counted equally, and converted occupancy to depth as *G*_i_ = −ln π_i_, with the deepest well in each condition set to zero. States never visited were assigned an upper bound on occupancy by the rule of three and reported as bounds rather than measurements. The two axes are unitless kinetic coordinates: states that exchange readily are placed close together and states that interconvert only through an intermediate are placed far apart, using multidimensional scaling of commute times from the pooled transition network so that every condition is drawn on the same layout. Because only six states exist, the continuous surface is an interpolation between them rather than a sampled density, with each state contributing a Gaussian well whose depth reproduces its measured G_i_, and each pair of connected states contributes a ridge at its midpoint whose height reflects the measured transition rate, *k*(i→j) = *P*(i→j) × sampling rate.

**Molecular dynamics simulations**

MspA’s structure was built by converting the atomic coordinates of its crystal structure (PDB: 1UUN)^4^ into the HyRes representation using the “at2hyres” script (<https://github.com/mdlab-um/HyRes_GPU>). To generate initial conformations of wild type and N29K/N30D p53-TAD, the disordered chains (residues 1-73) was built using CHARMM^5,6^. Generation of 300 conformations of both variants was carried out by simulating each chain at 400 K for 50 ns. Simulations were performed with the HyRes coarse-grained protein model^7^ in CHARMM interfaced to OpenMM^8^ on the CUDA platform (mixed precision). Solvent was treated implicitly with Debye–Hückel electrostatic screening at an ionic strength of 0.15 M. Nonbonded interactions used atom-based switching, with van der Waals interactions switched over 16–18 Å and a 20 Å nonbonded list cutoff. Each system was minimized by 500 steps of steepest descent followed by 500 steps of adopted-basis Newton–Raphson, then propagated by Langevin dynamics (leapfrog integrator, 4 fs timestep) with a friction coefficient of 0.2 ps⁻¹ on non-hydrogen atoms; bonds to hydrogen were constrained using SHAKE (tolerance 10⁻⁶). Coordinates were saved every 6 ps. Following generation of the 300 starting conformations of each p53-TAD variant, individual steered MD systems were built with MspA’s COM at the origin (cis opening pointing in +Z direction) and p53-TAD conformations’ COM placed 5 nm directly above MspA’s opening. The MspA pore was held near its reference geometry by harmonic positional restraints (k = 1.0 kcal mol⁻¹ Å⁻²) applied to all pore Cα atoms. Translocation was driven by a constant external force applied along the −Z (pore) axis, distributed equally over the acidic Cα sites (Asp and Glu) of p53-TAD, for a total pulling force of 10 pN across the protein. To prevent lateral escape of the domain from the pore region before capture, a half-harmonic cylindrical restraint (MMFP GEO in CHARMM) was imposed on p53-TAD’s Cα atoms about the Z axis: the potential was zero for radial distances r ≤ 30 Å² and harmonic beyond, U(r) = ½·k·(r − 30)² for r > 30 Å, with k = 5 kcal mol⁻¹ Å⁻². Pulling simulations were run at 300K for 200ns each. For the bulk simulations, 4 conformations of each variant were minimized by 500 steps of steepest descent followed by 500 steps of adopted-basis Newton–Raphson, and subsequently simulated for 400ns under the same conditions as the steered MD (minus the external force and cylindrical restraint).

**Simulation analysis**

Analyses were performed on 300 pulling trajectories per construct using MDAnalysis^9^. All coordinates were referenced to the center of mass of the MspA constriction (chains MSP1–MSP8, residues 90–93, Cα), so Z = 0 marks the constriction, +Z the cis vestibule, and R the radial distance from the pore axis; the pore cross-section shown in figures was obtained per Z bin from the minimum and maximum atomic radii of MspA, padded by 1.7 Å and boxcar-smoothed over 5 bins. p53-TAD occupancy was quantified from aggregate 2D (R, Z) histograms of the TAD center of mass normalized to a joint density, with the marginal P(Z) obtained by integration and PMF(Z) =$-\ln\frac{P(Z)}{P_{max}}$ in $k_{B}T$. Construct differences were taken as wild type minus N29K/N30D, with ΔPMF(Z) computed under a common normalization and summarized per Z region as the mean ($k_{B}T$). Conformational analysis was performed within Z regions of 0–30 Å (near constriction) and 35–70 Å (vestibule). Cα coordinates were sampled every 10th frame, cached into 5 Å Z slabs, and re-binned by instantaneous center-of-mass Z. Each frame was represented by the upper triangle of its Cα–Cα distance matrix (a rotation- and translation-invariant descriptor) and standardized/reduced by PCA retaining 90% of the variance (≤ 25 components). Because this descriptor encodes chain shape only and carries no axial coordinate, frames from different Z regions are directly comparable. For visualization, frames from both regions and both constructs were reduced together and projected to two dimensions by UMAP. Clustering used a separate reduction per Z region, with wild type and N29K/N30D pooled within the region, partitioning frames by Gaussian-mixture modeling (full covariance) into two states per region. This ceiling was used deliberately to yield a small number of well-separated and interpretable representative conformations, as configuration space in the vestibule would otherwise yield many clusters with similar characteristics to one another. Each cluster was represented by its medoid (the member closest to the cluster centroid in that region's PCA space). Ensembles were visualized by overlaying the 25 frames nearest each medoid in its region's feature with overlaid chains rotated about Z onto the medoid by 2D orthogonal Procrustes alignment.

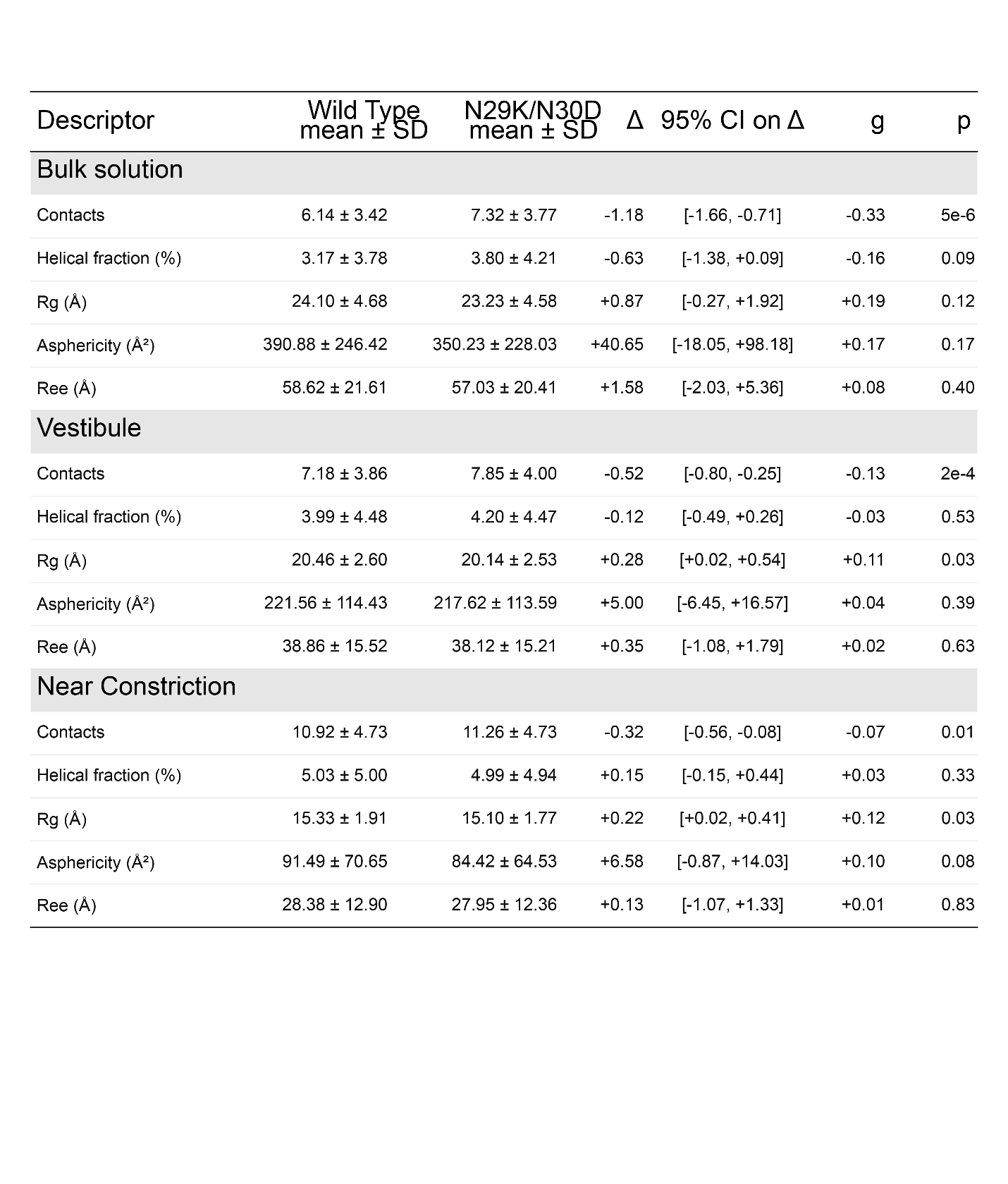

**Table S1: Simulated conformational descriptors of p53.** Frame-level mean ± SD per construct, the difference Δ = wild type − N29K/N30D (positive = higher in wild type), its 95% run-level bootstrap CI, Hedges' *g*, and the bootstrap *p* value. Helical fraction is given as a percentage of residues.

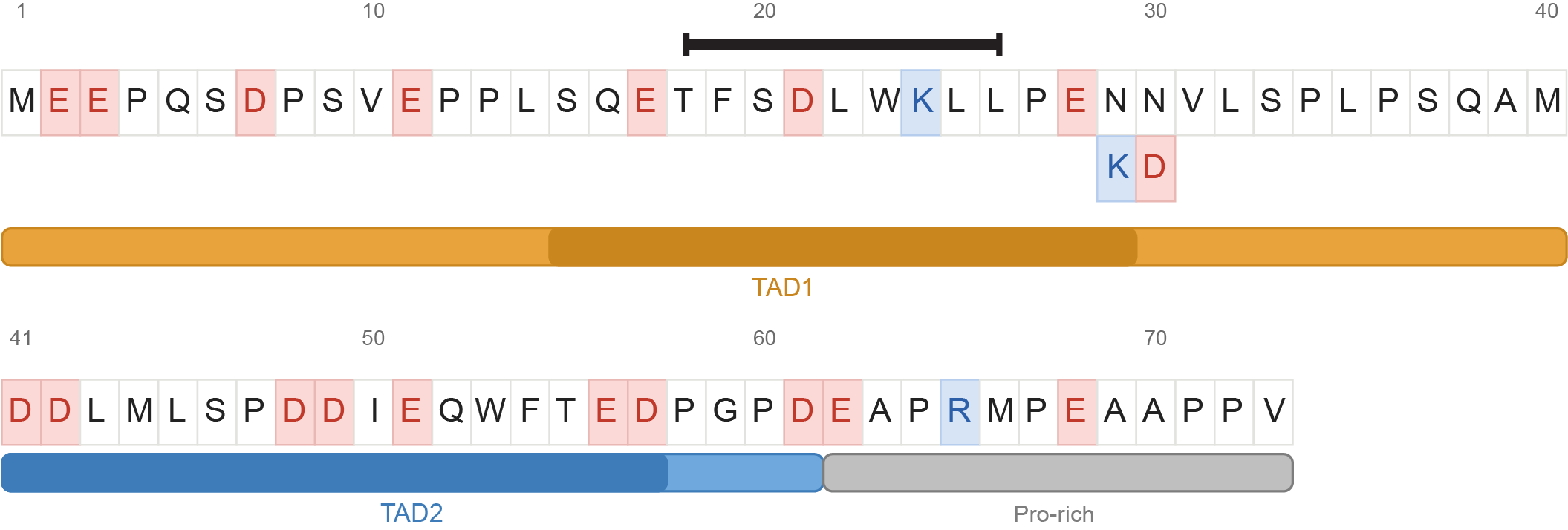

**Figure S1: p53-TAD primary sequence annotated.** Primary sequence of p53’s transactivation domain colored based on charge (red: acidic, blue: basic, black: uncharged). Names of the subdomains of p53-TAD are shown below the primary sequence. Black bar above residues 18-26 represents the binding site of MDM2 and the K/D below residues 29/30 shows the cancer-associated mutational site.

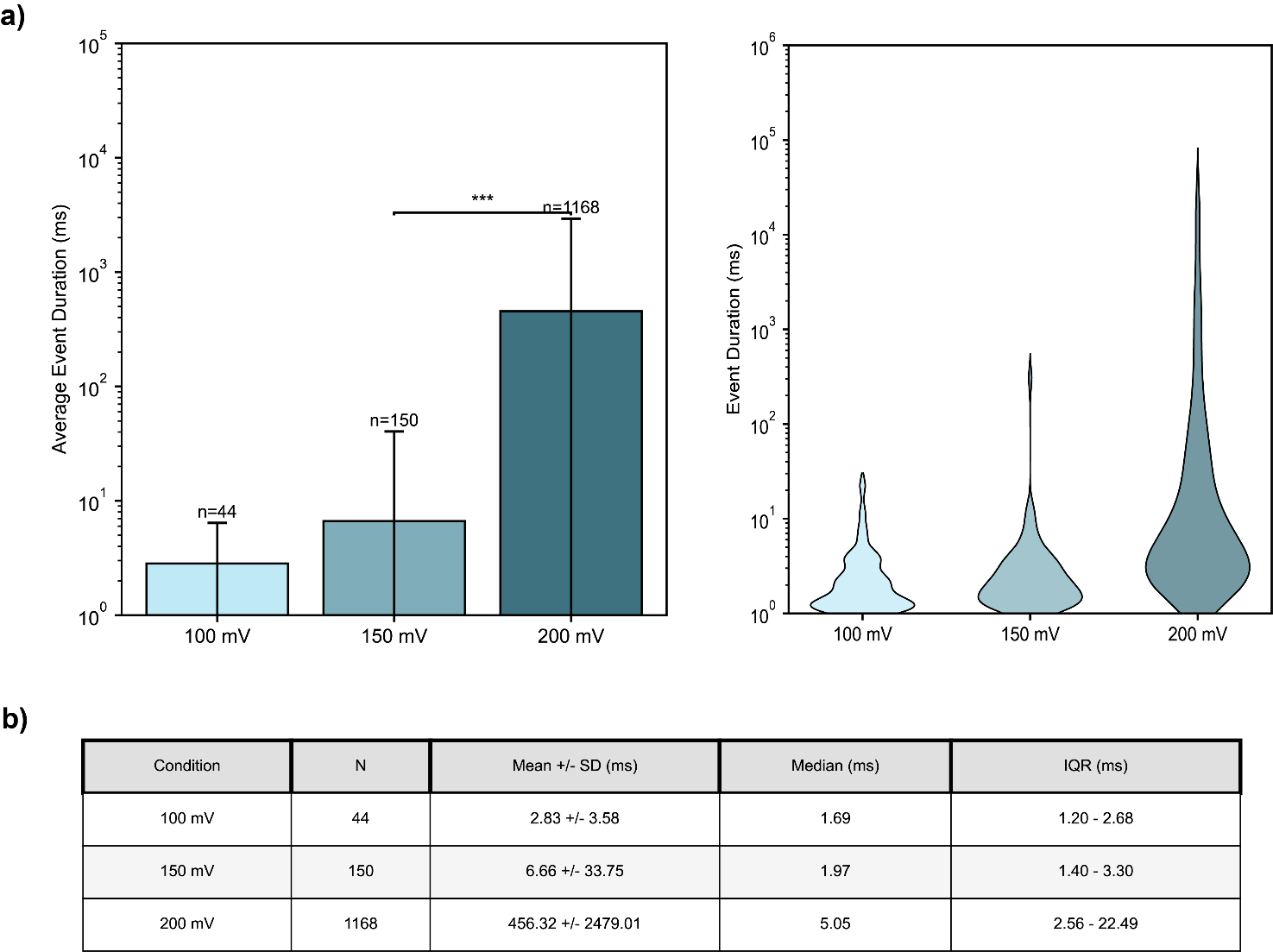

**Figure S2: Duration of wild type p53-TAD voltage study events. a)** Bar chart showing the average event duration ± standard deviation (left) and violin plot (right) of wild type p53-TAD events at different voltages. **b)** Table displaying duration statistics of wild type p53-TAD voltage events.

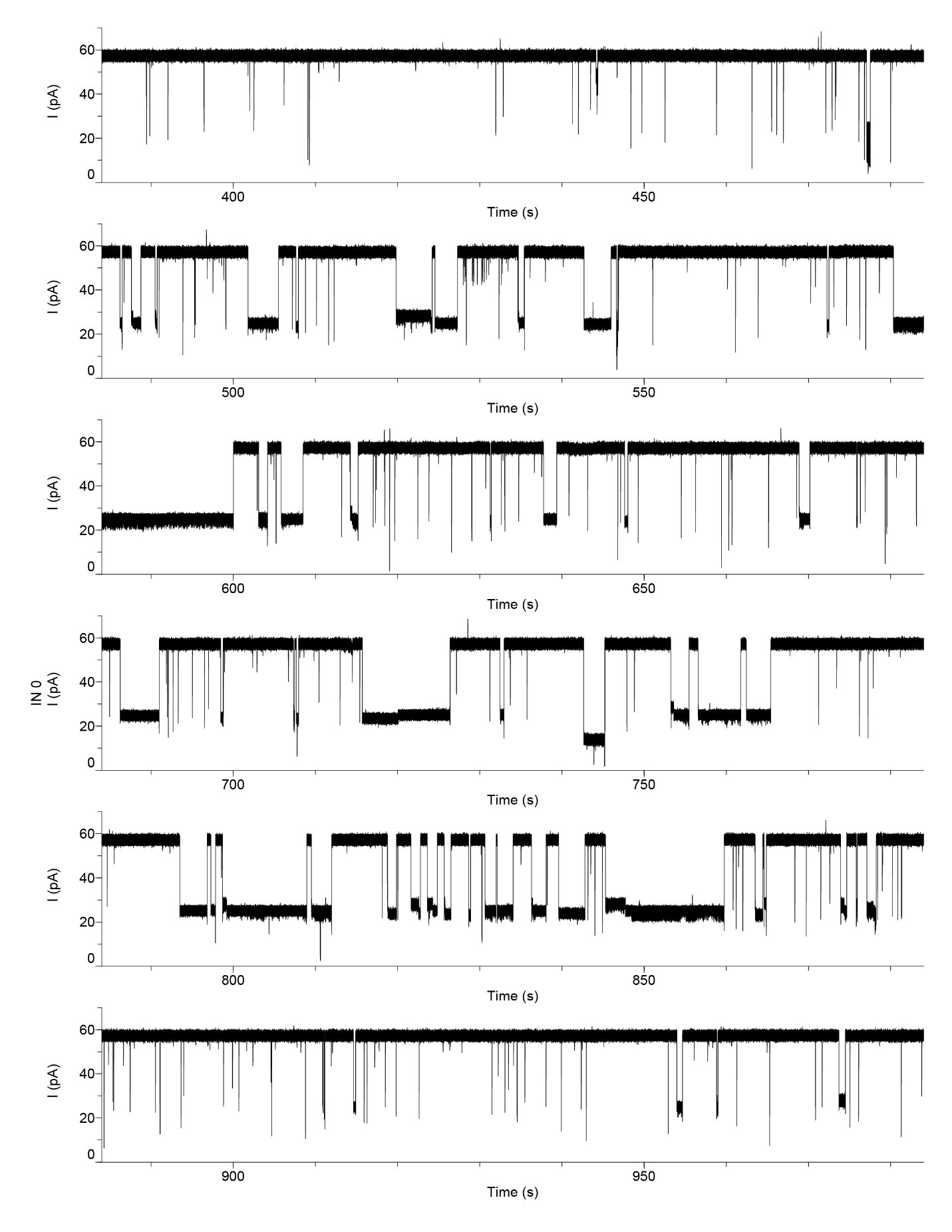

**Figure S2: Representative long current trace of wild type p53-TAD capture under +200mV.** Representative 10 min trace of wild type p53-TAD capture in 150mM NaCl, 20mM HEPES, pH 7.4 solution under +200mV applied voltage.

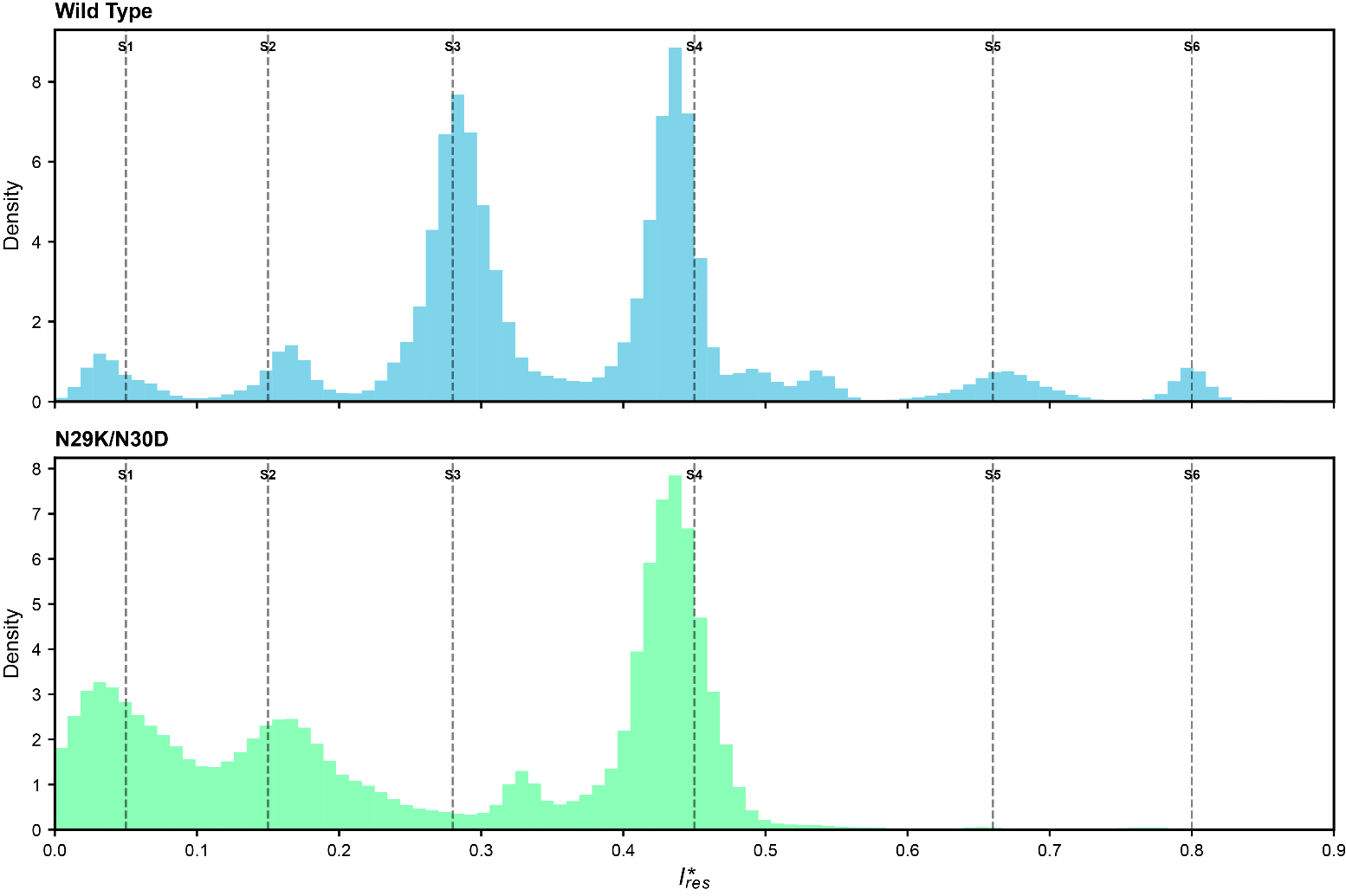

**Figure S4: All points histograms of wild type and N29K/N30D +200mV events**. All points histograms (100 bins) of all wild type (blue/top) and N29K/N30D (green/bottom) capture events. State assignment means notated as dashed lines above the $I_{res}^{*}$ peaks corresponding to that particular state.

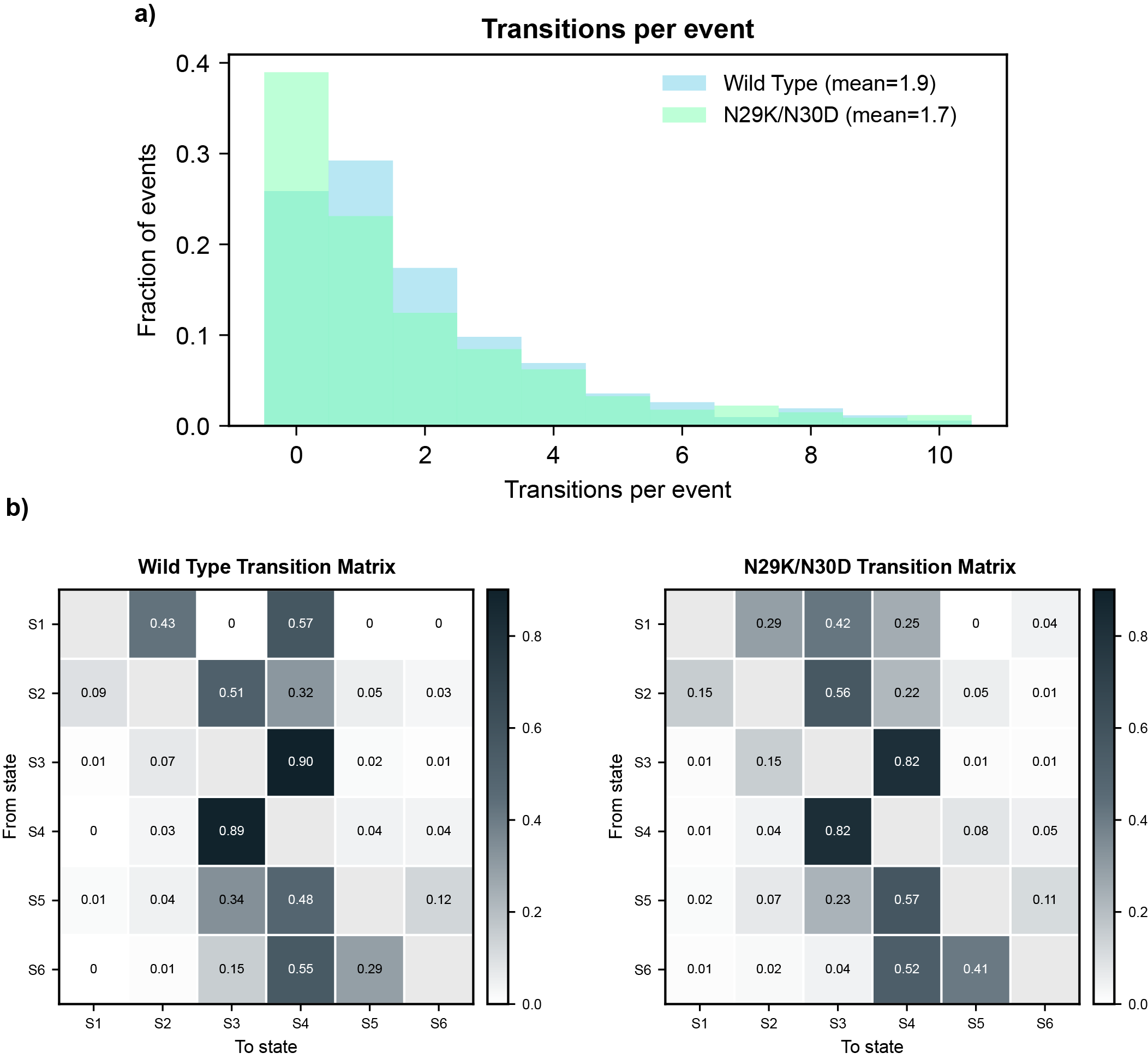

**Figure S5: State transitions per event and transition matrices. a)** Histogram displaying the distribution of transitions per event of wild type (blue) and N29K/N30D (green) p53-TAD capture events. **b)** Off-diagonal normalized transition matrices of wild type (left) and N29K/N30D capture events demonstrating the probability of exiting from one state to the next.

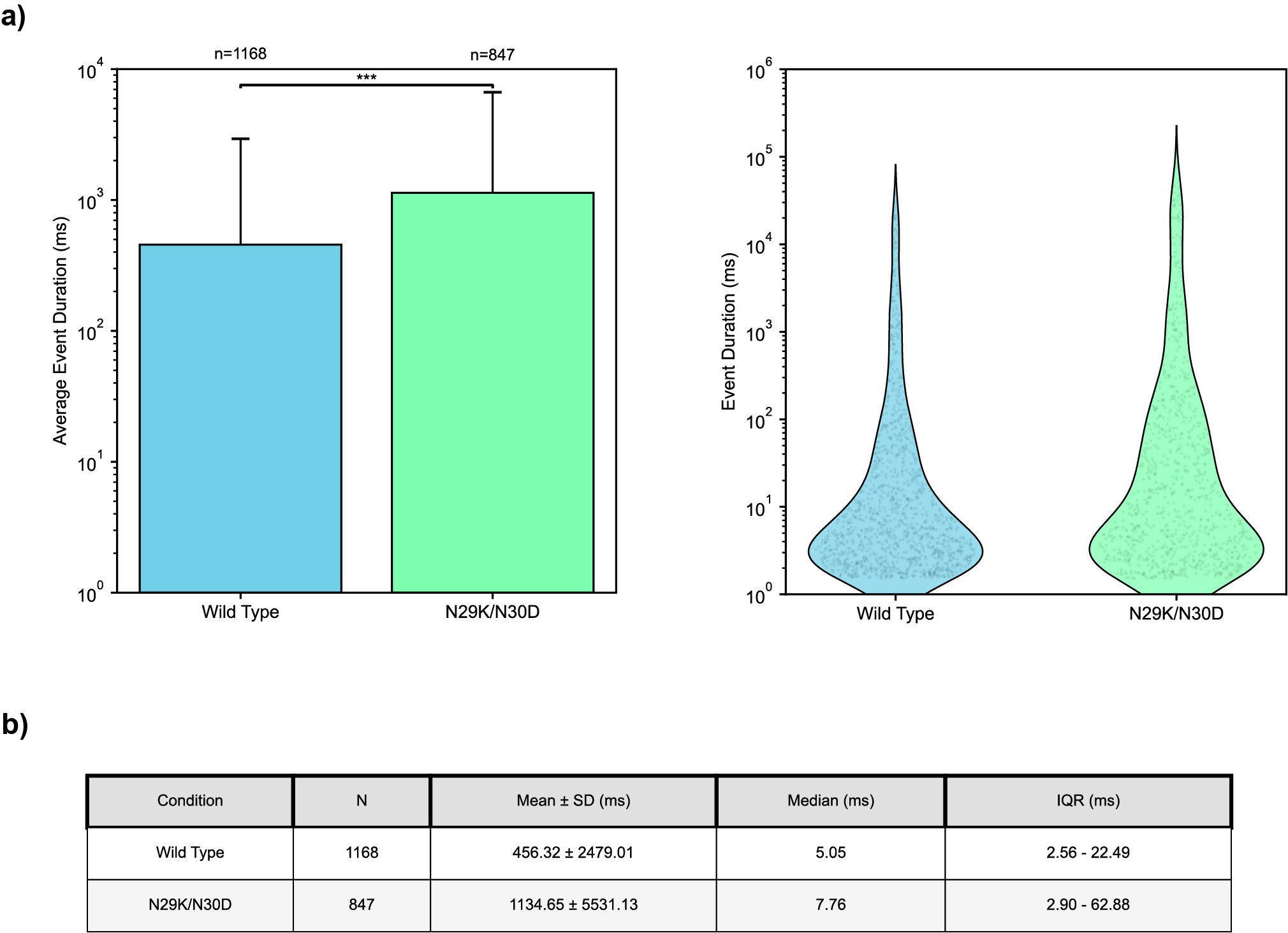

**Figure S6: Duration of wild type and N29K/N30D p53-TAD +200mV events. a)** Bar chart showing the average event duration ± standard deviation (left) and violin plot (right) of WT and N29K/N30D +200mV events. **b)** Table displaying duration statistics of WT and N29K/N30D +200mV events.

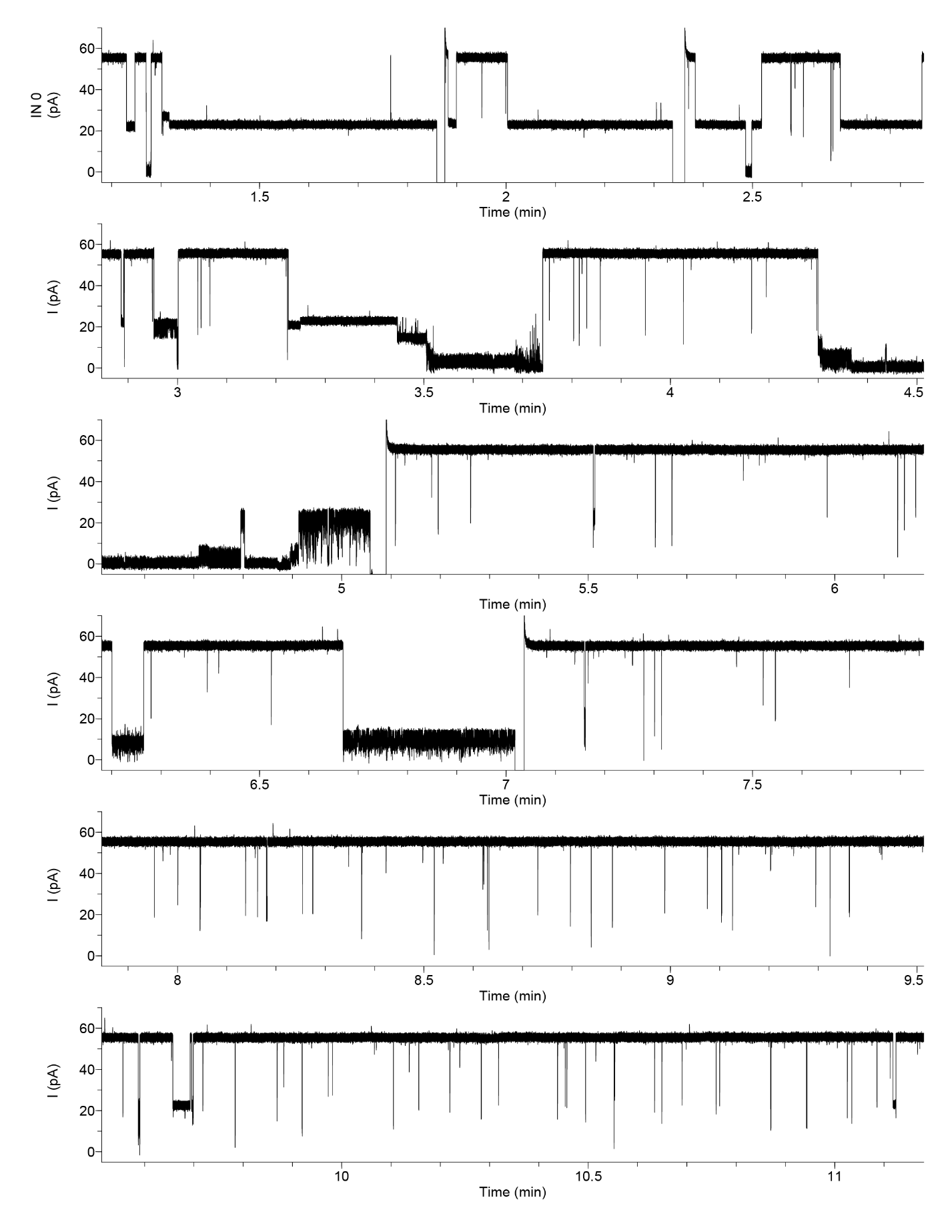

**Figure S7: Representative long current trace of N29K/N30D p53-TAD capture under +200mV.** Representative 10 min trace of N29K/N30D p53-TAD capture in 150mM NaCl, 20mM HEPES, pH 7.4 solution under +200mV applied voltage.

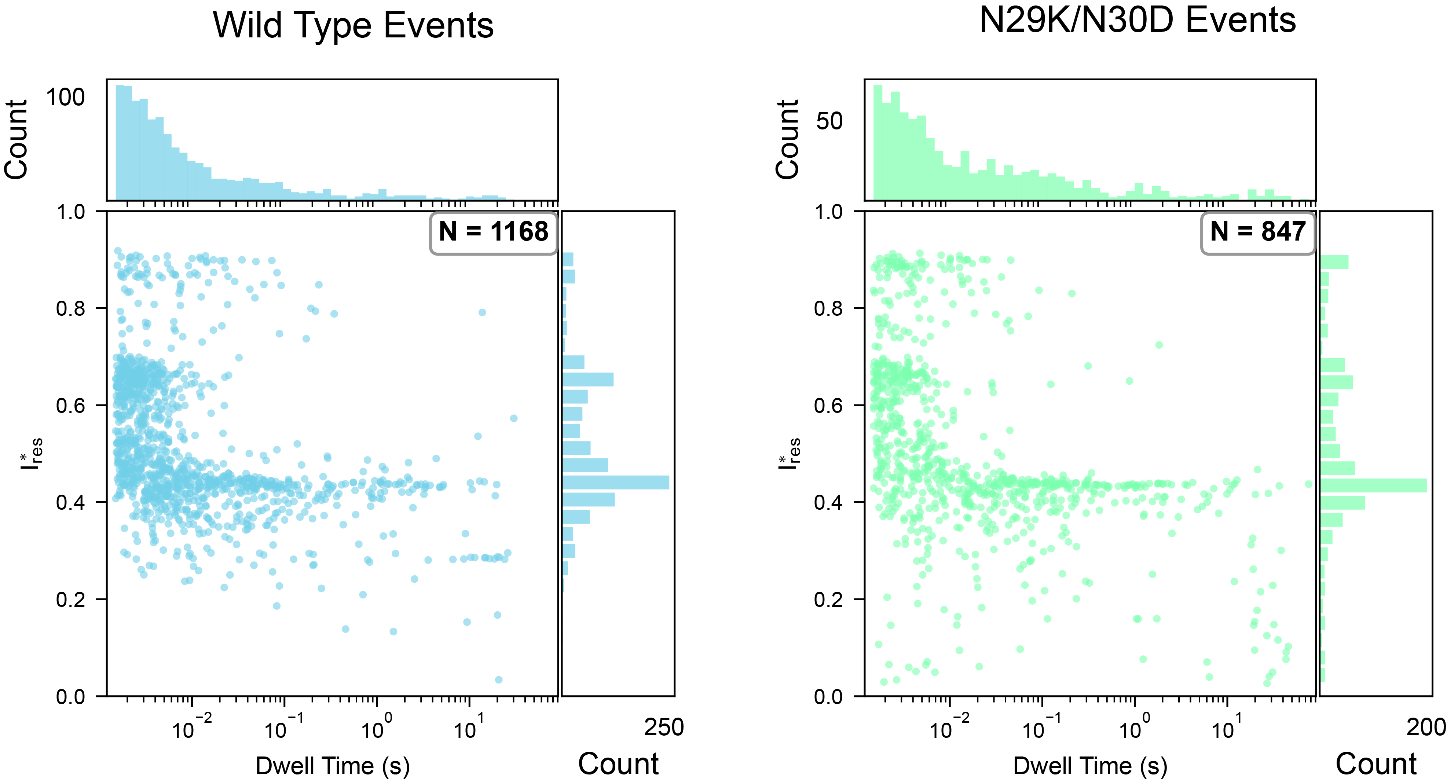

**Figure S8: Blockage vs duration scatterplots of wild type and N29K/N30D p53-TAD capture events.** Individual events’ average $I_{res}^{*}$ is plotted against dwell time for wild type (left/blue) and N29K/N30D (right/green) p53-TAD capture under +200mV. Marginal distributions for $I_{res}^{*}$ (25 bins) and Dwell time (50 bins) displayed to the right and top of the plots respectively. N indicates number of total events plotted.

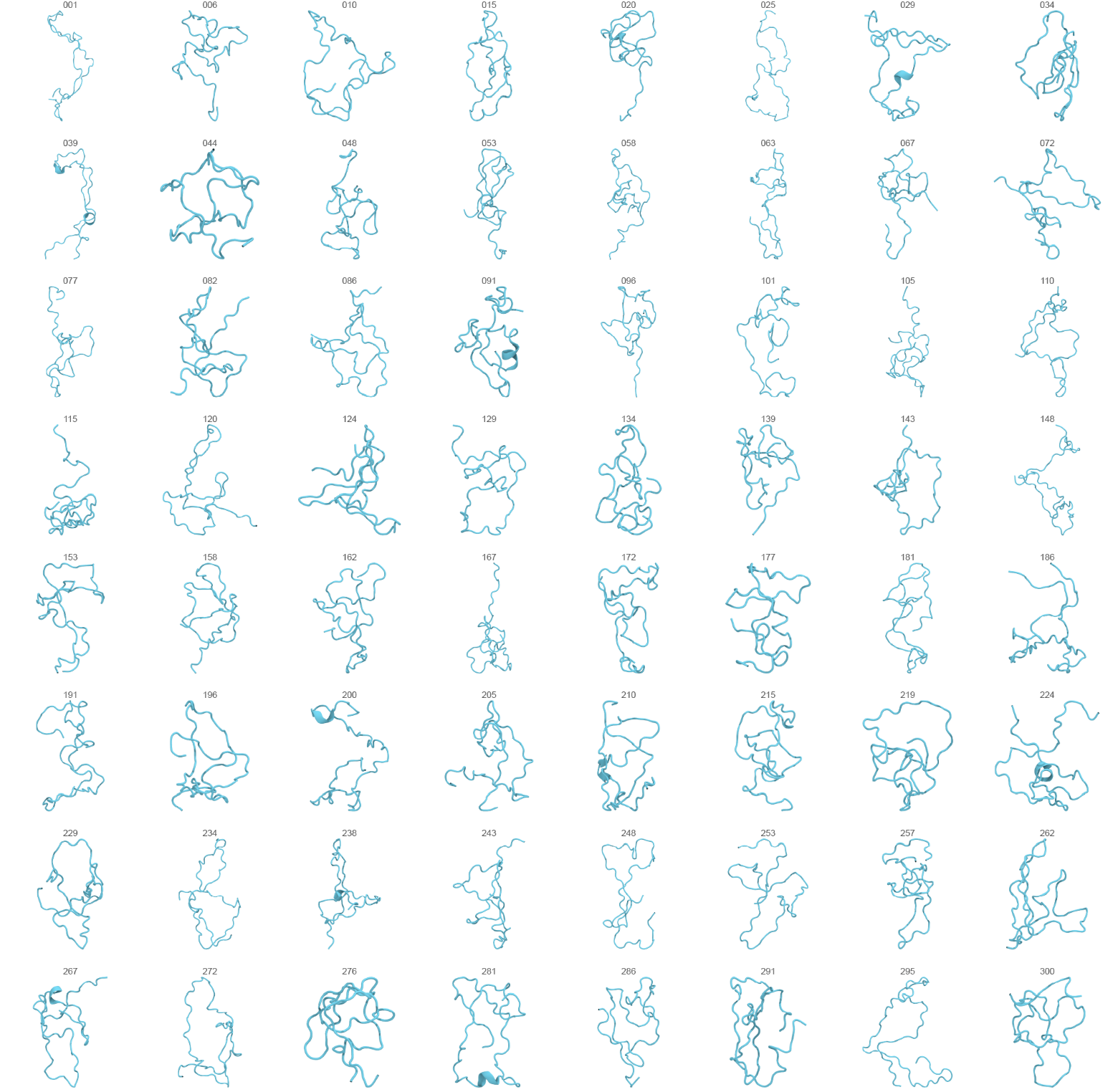

**Figure S9: Representative starting conformations of wild type p53-TAD for SMD.** Shown is 64 of the 300 starting conformations of wild type p53-TAD used at the start of each steered MD simulation. Number above structure represents the simulation number used.

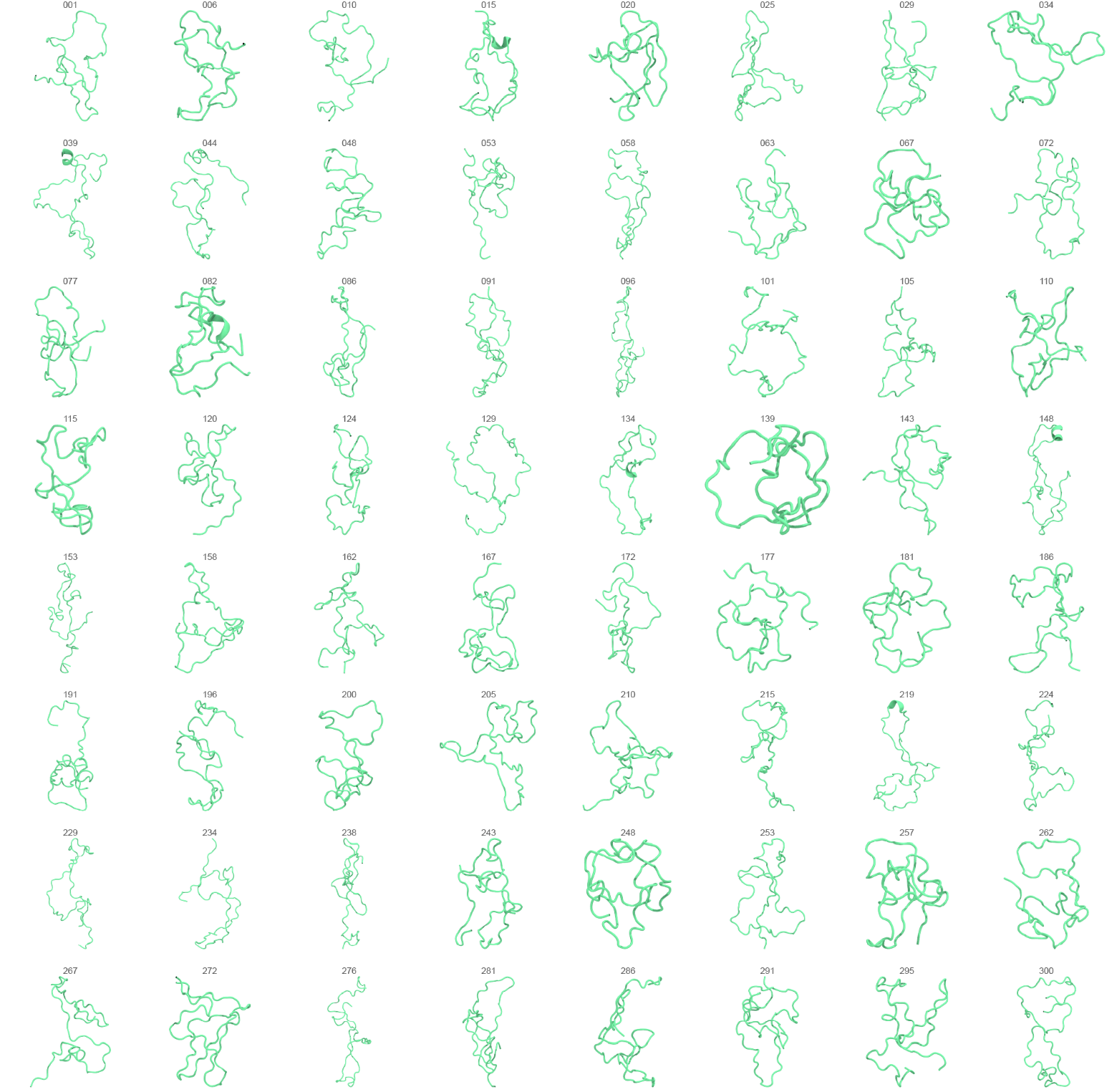

**Figure S10: Representative starting conformations of N29K/N30D p53-TAD for SMD.** Shown is 64 of the 300 starting conformations of N29K/N30D p53-TAD used at the start of each steered MD simulation. Number above structure represents the simulation number used.

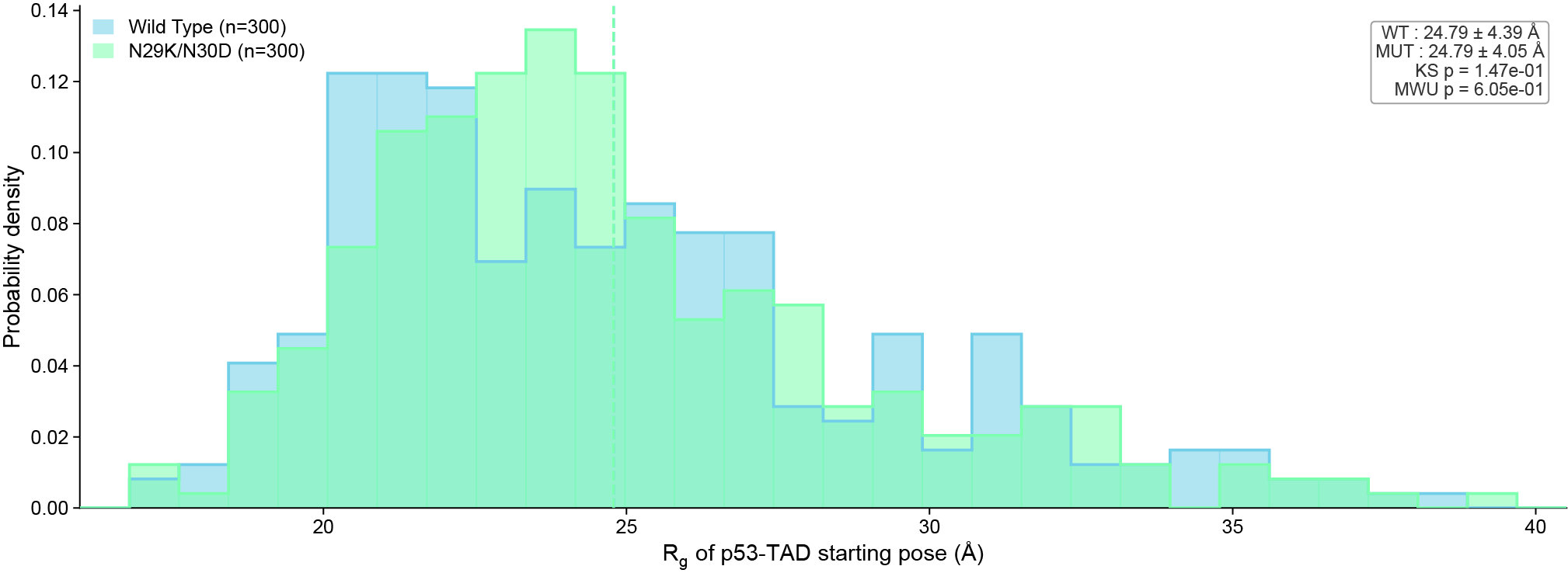

**Figure S11: Radius of gyration of starting SMD conformational ensembles.** Overlapping histograms (30bins) of the radius of gyration of the starting conformations of WT and N29K/N30D p53-TAD used in SMD (vertical dashed lines represent average). Upper right legend reports the mean for both WT and N29K/N30D as well as the Kolmogorov-Smirnov (KS) and Mann-Whitney (MW) p-values.

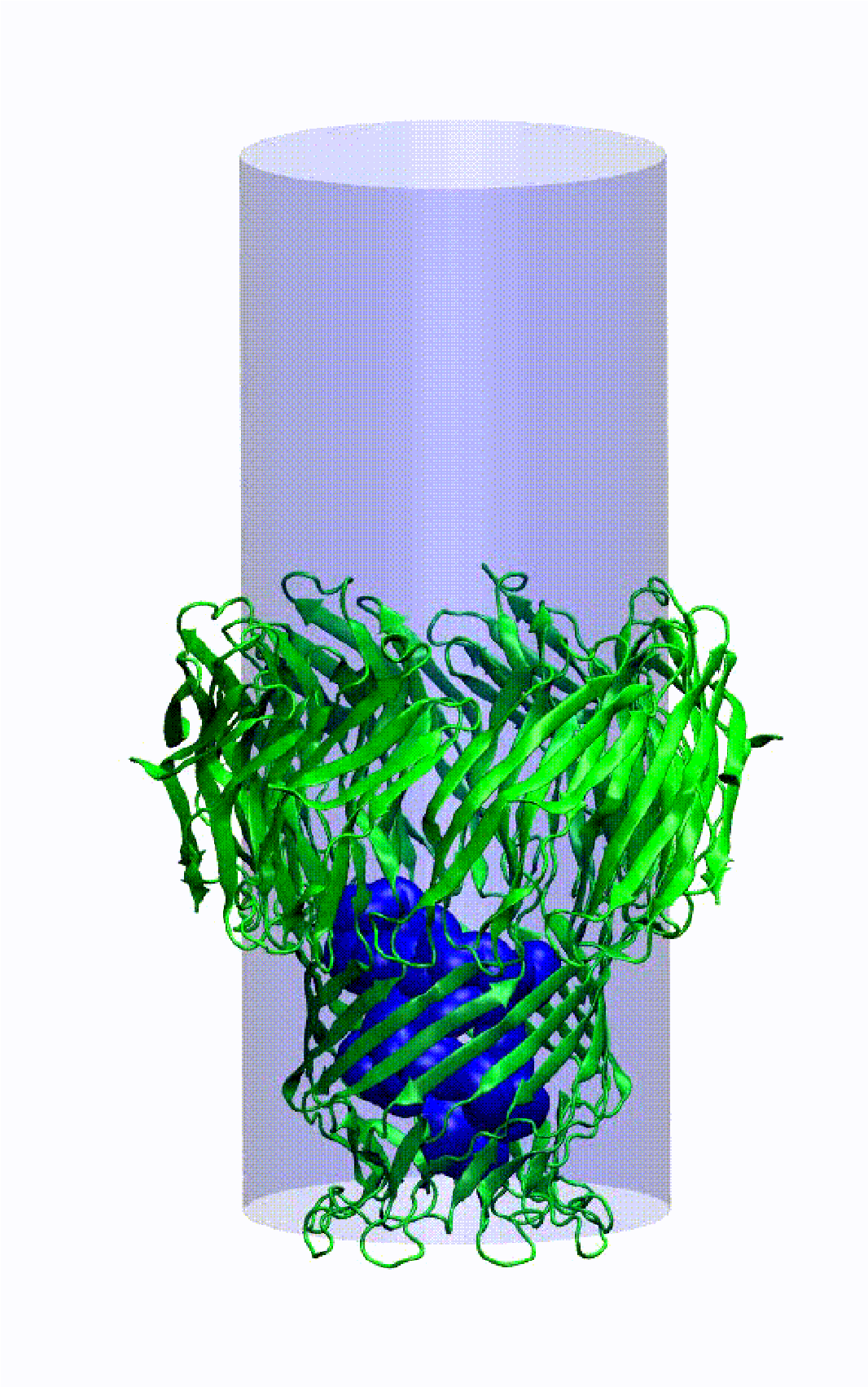

**Figure S12: Simulation system for SMD.** p53-TAD (dark blue) is pulled into MspA (green) via 10pN constant force in the -Z direction (cis to trans) with an additional cylindrical potential (blue cylinder) pushing any Ca of p53-TAD that crosses it back into the center of the cylinder.

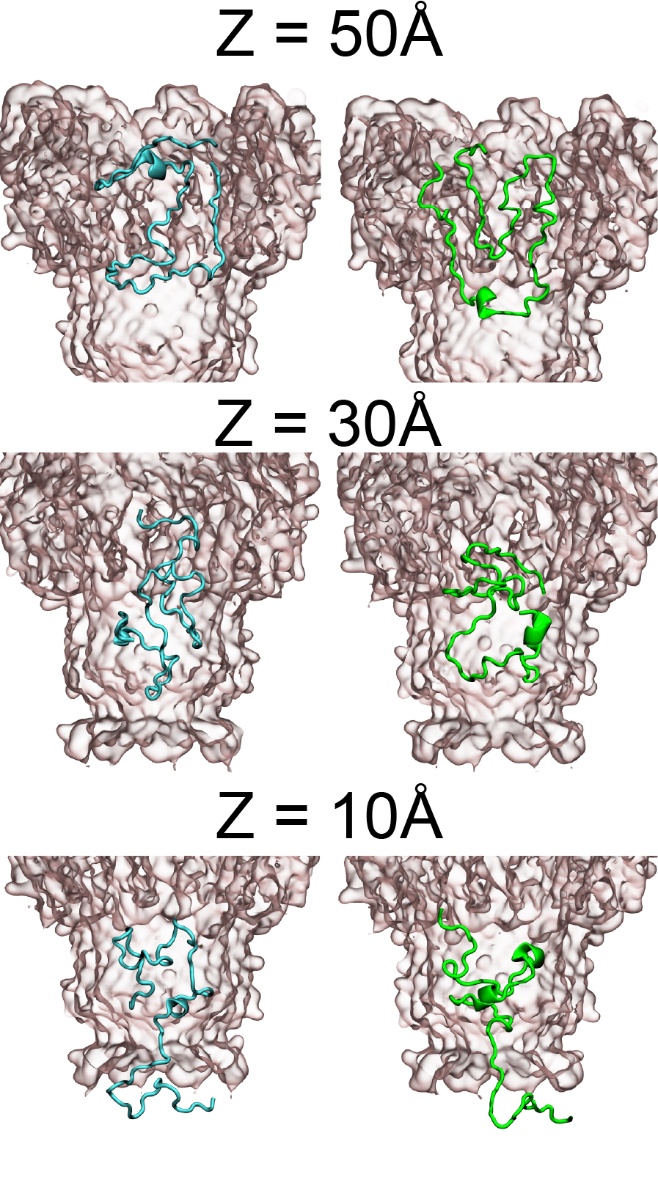

**Figure S13: Representative snapshots of p53-TAD at occupied Z positions.** p53-TAD (dark blue) is pulled into MspA (green) via 10pN constant force in the -Z direction (cis to trans) with an additional cylindrical potential (blue cylinder) pushing any Ca of p53-TAD that crosses it back into the center of the cylinder.

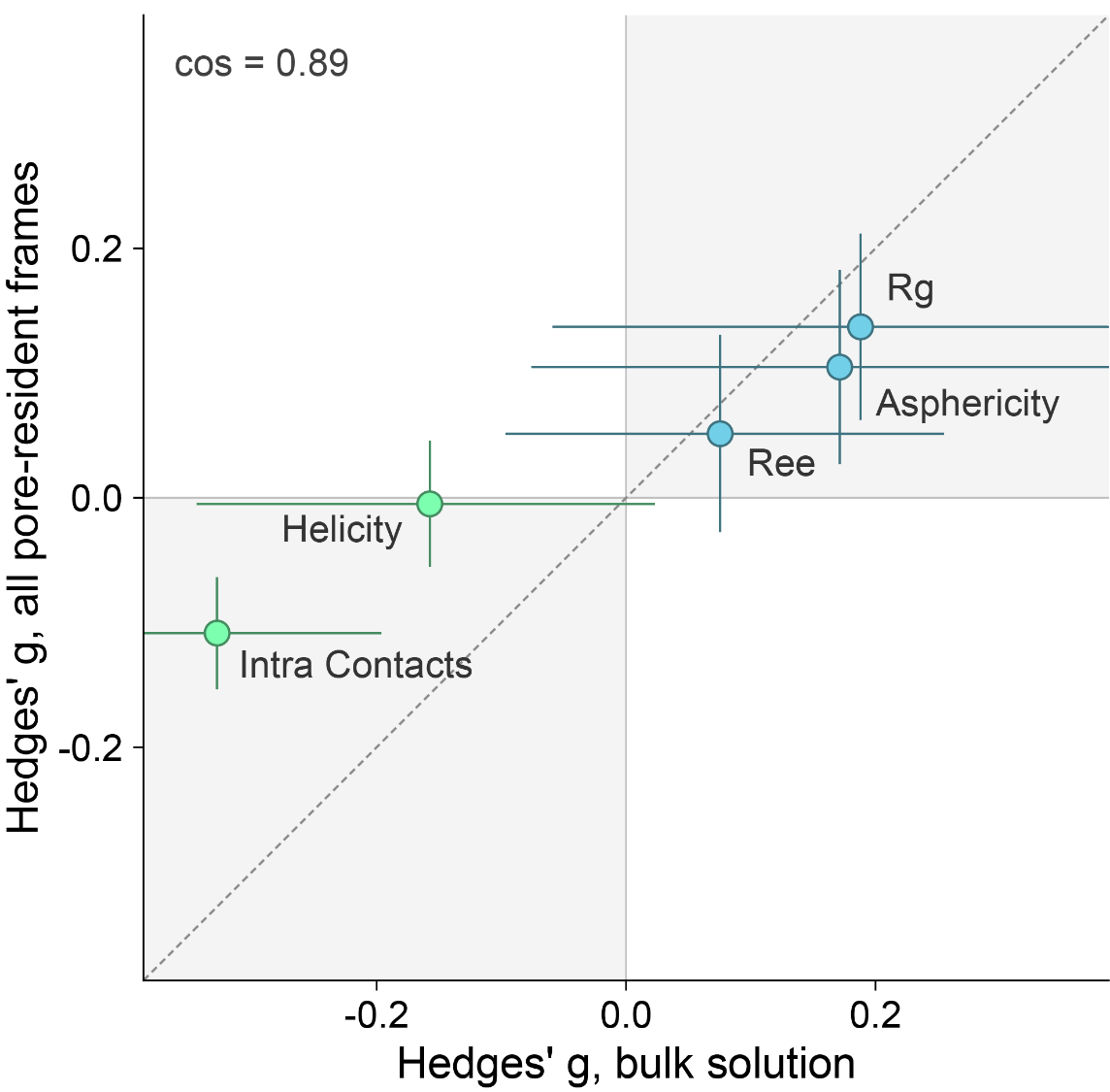

**Figure S14: Mutational effect on p53-TAD conformation is preserved under confinement.** Hedges' g for WT − N29K/N30D in bulk solution against pore-resident frames with one point per descriptor. Bars are 95% run-level bootstrap CIs.

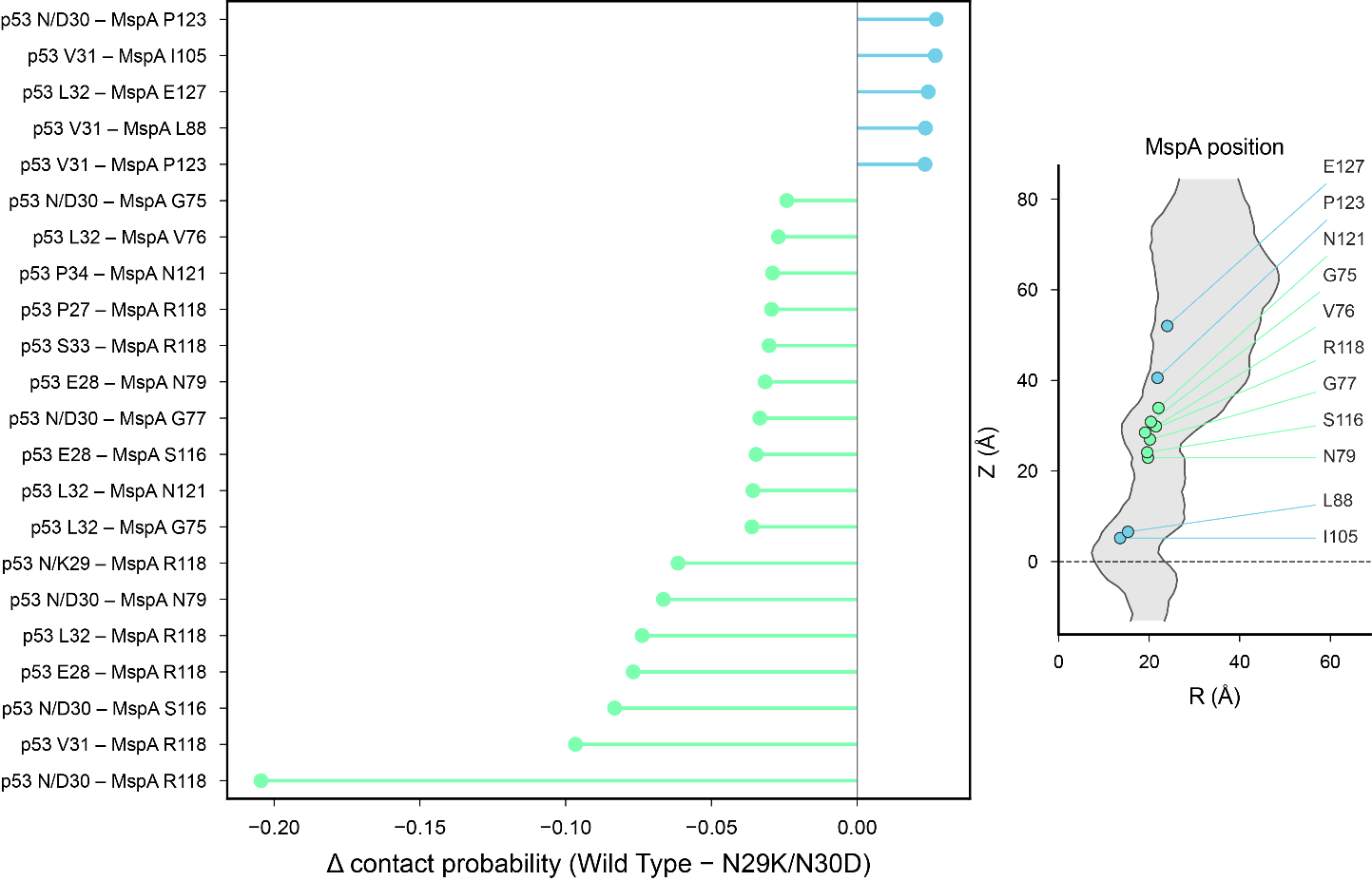

**Figure S15: Top contact difference between wild type and N29K/N30D p53-TAD and MspA.** Difference in contact probability between p53-TAD and MspA shown on left (green and negative: enriched in mutant, blue and positive: enriched in wild type). Right displays the location of the top differential contact spots of MspA.

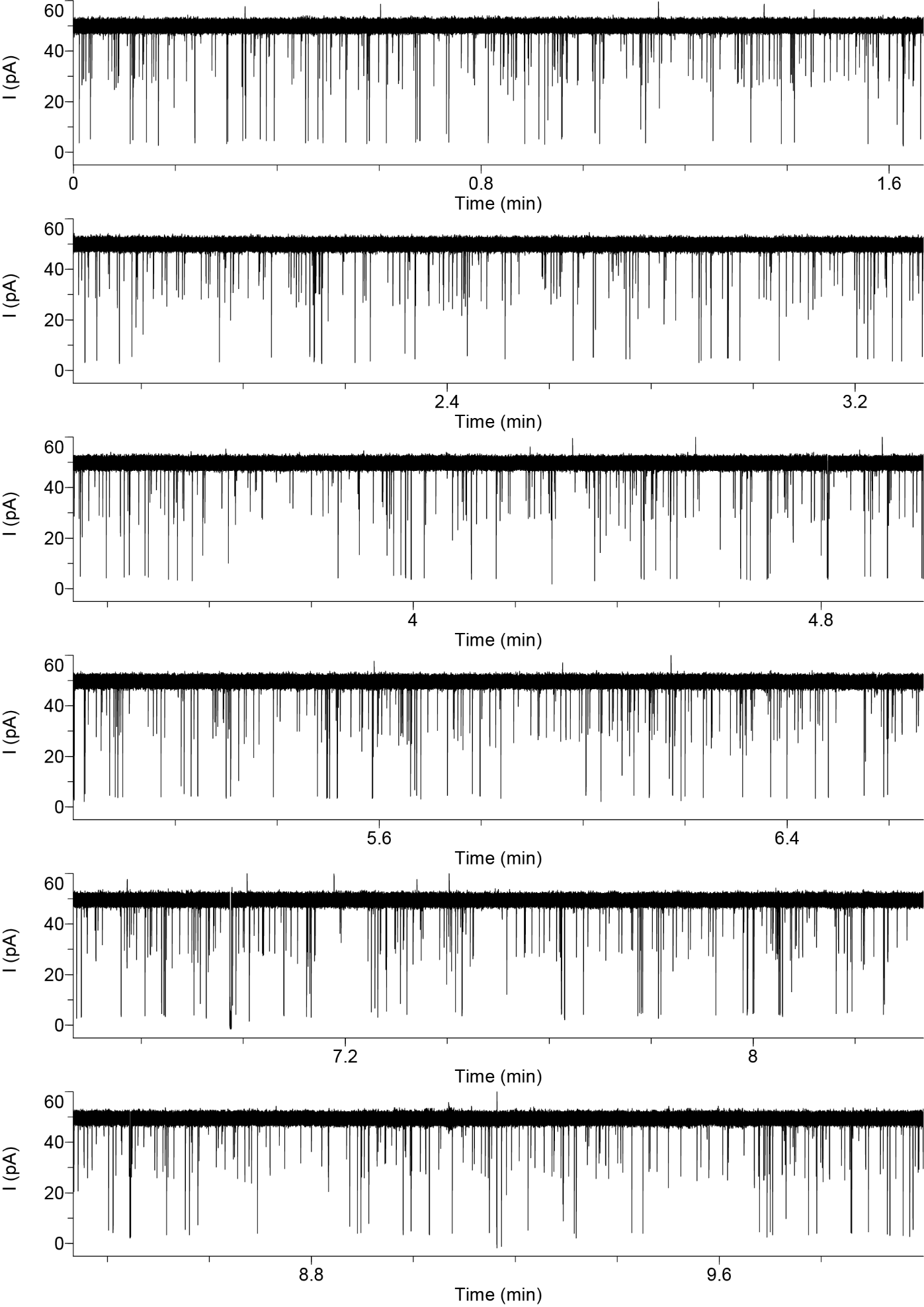

**Figure S16: Representative long current trace of MDM2 Control under +200mV.** 10minute trace of 400nM MDM2 events under +200mV applied potential.

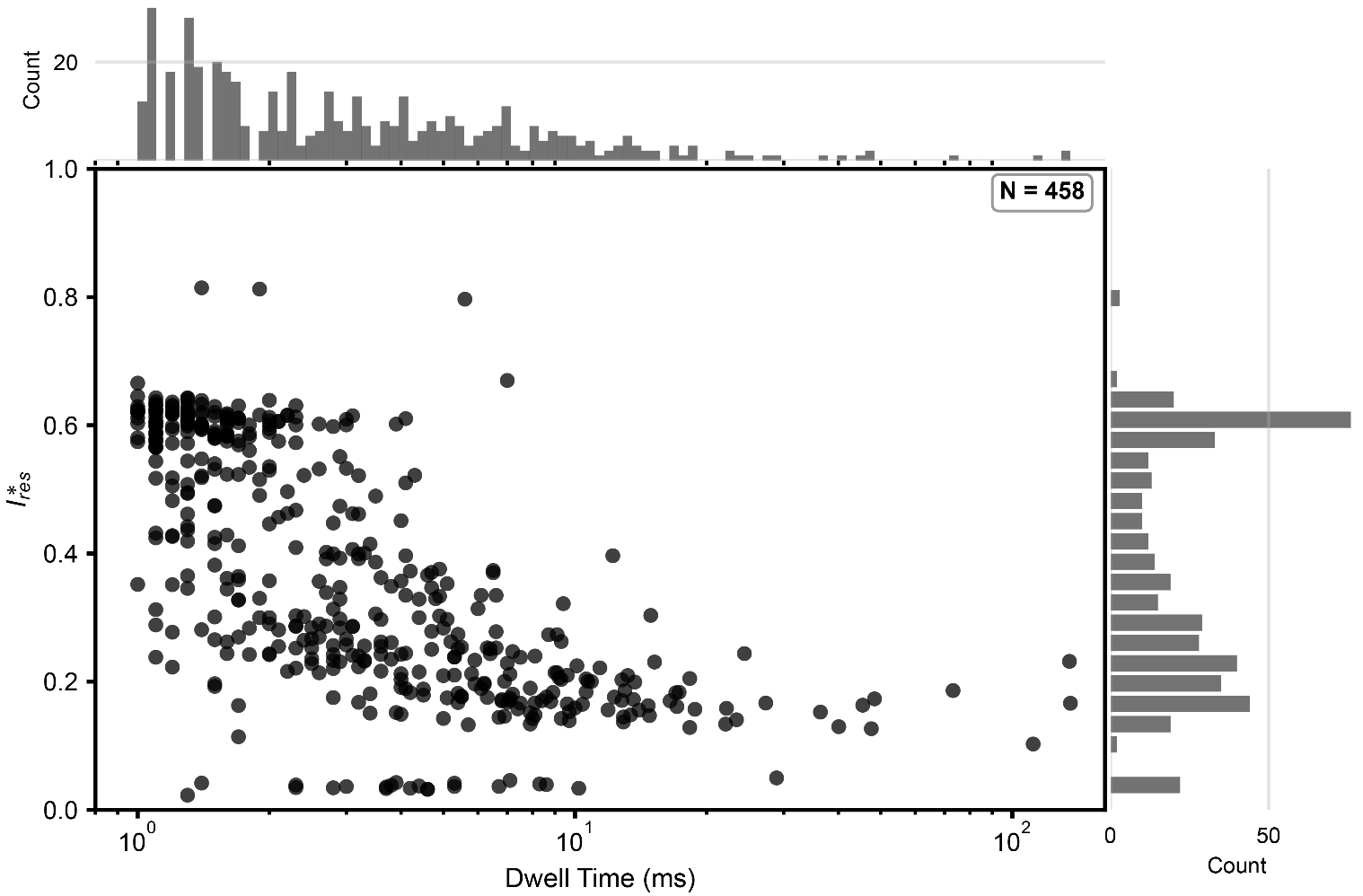

**Figure S17:** $\boldsymbol{I}_{\boldsymbol{res}}^{\boldsymbol{*}}$ **vs dwell time for MDM2 control events.** Individual MDM2 control events plotted as a function of duration (x axis) vs conductance blockage (y axis). Marginal distributions for x (100 bins) and y (25 bins) shown above and to the right of the plot respectively.
